# Crumbs2 mediates ventricular layer remodelling to form the adult spinal cord central canal

**DOI:** 10.1101/744888

**Authors:** Christine Tait, Kavitha Chinnaiya, Mariyam Murtaza, John-Paul Ashton, Nicholas Furley, Chris J. Hill, C Henrique Alves, Jan Wijnholds, Kai S. Erdmann, Raman M. Das, Penny Rashbass, Kate G. Storey, Marysia Placzek

**Affiliations:** Department of Biomedical Science and Bateson Centre, University of Sheffield, UK; Department of Ophthalmology, Leiden University Medical Centre, Netherlands Institute for Neuroscience, The Netherlands; Division of Cell & Developmental Biology, School of Life Sciences, University of Dundee, UK

**Author notes:** Department of Integrative Medical Biology, Umea University, Sweden (Christine Tait). Division of Molecular and Cellular Function, University of Manchester, UK. Equal contribution (Christine Tait, Kavitha Chinnaiya).

## Abstract

In the spinal cord, the adult central canal forms through a poorly-understood process termed dorsal collapse that involves attrition and remodelling of the pseudostratified dorsal ventricular layer. Here we show, in mouse, that dorsal ventricular layer cells adjacent to midline Nestin^(+)^ radial glia downregulate the apical polarity proteins Crumbs2 (CRB2) and aPKC and delaminate in a step-wise manner; concomitantly, Nestin^(+)^ radial glial end-feet ratchet down, to repeat this process. Nestin^(+)^ radial glia secrete a factor that promotes cell delamination. This activity is mimicked by a secreted variant of CRB2 (CRB2S), which is specifically expressed by dorsal midline Nestin^(+)^ radial glia. In cultured cells, CRB2S associated with apical membranes and decreased cell cohesion. Analysis of *Crb2^F/F^/Nestin-Cre^+/−^* mice further confirmed an essential role for CRB2 in dorsal collapse. We propose a model in which CRB2S promotes the progressive attrition of the ventricular layer without loss of overall integrity. This novel mechanism may operate more widely to promote orderly progenitor delamination.

## Introduction

The ventricular layer (VL) of the embryonic spinal cord is composed of pseudostratified radial glial stem/progenitor cells that line the central lumen. VL cells express SoxB1 proteins (Canizares et al., 2019), a feature of their neuroepithelial origin (reviewed in Gouti et al., 2015; Pevny and Placzek, 2005), and differentially express homeodomain transcription factors, a feature of dorso-ventral patterning (reviewed in Jessell, 2000; Le Dreau and Marti, 2012; Ulloa and Briscoe, 2007). In early embryogenesis, VL cells undergo neurogenesis in a process that involves apical constriction, adherens-junction disassembly, acto-myosin-mediated abscission and medio-lateral migration (Das and Storey, 2014; Kasioulis et al., 2017). Following the major period of neurogenesis, VL cells switch to gliogenesis, and glial cells migrate out of this layer (Deneen et al., 2006; Stolt et al., 2003; Hochstim et al., 2008) reviewed in (Laug et al., 2018).

Concomitant with the transition to gliogenesis (around E12 in the mouse) the VL begins to remodel, to ultimately give rise to the ependymal layer of the adult central canal. Ependymal cells constitute key components of a quiescent stem cell niche (Johansson et al., 1999; Sabourin et al, 2009; reviewed in Marichal et al., 2017), are implicated in glial scar formation after spinal injury (Meletis et al., 2008; Li et al., 2016; reviewed in Marichal et al., 2017), and serve important mechanical and sensory functions (reviewed in Bruni, 1998; Del Bigio, 1995). Multiple steps contribute to the remodelling of the VL into the ependymal layer (Cañizares et al, 2019; Xing et al., 2018), including dorsal collapse, a delamination of dorsal VL cells that results in a pronounced dorso-ventral reduction in the length of the lumen (Altman and Bayer 1984; Böhme, 1988; Fu et al., 2003; Sevc et al., 2009; Shinozuka et al., 2019; Spassky et al., 2005; Sturrock, 1981; Yu et al., 2013; Xing et al., 2018). Little is understood, however, of the mechanisms that mediate dorsal collapse.

Proteins of the Crumbs (Crb) and Par complexes (the latter composed of PAR3/Par6/aPKC) are present on the apical side of epithelial cells. In invertebrates, Par and Crumbs proteins directly interact to determine the apico-basal axis and the position and stability of cell-cell adherens junctions (Wodarz et al., 1995; Klebes and Knust, 2000; Goldstein and Macara, 2007; Assemat et al;., 2008; Bulgakova and Knust, 2009; reviewed in Pichaud, 2019). Par and Crumbs complex components are evolutionarily conserved, and similarly regulate polarity and integrity of vertebrate epithelia and neuroepithelia (reviewed in McCaffrey and Macara, 2011; Nikolopoulou et al., 2018); in the mouse telencephalon, CRB2 is required for maintenance of the apical polarity complex (Dudok et al., 2016) and in zebrafish, Crb2a controls the localization of tight and adherens junction proteins in cardiomyocytes (Jiménez-Amilburu and Stainier et al., 2019). Apical complexes play a central role in early stages of neural tube formation (reviewed in Cearns et al., 2016). Moreover, the finding in zebrafish that mutation in *pard6yb* results in the failure of dorsal collapse (Kondrychyn et al 2008; Munson et al., 2008) suggests that apical polarity complex regulation plays a critical role in VL remodelling. However, it remains unclear whether and how components of the apical polarity complex change during dorsal collapse, nor in which cell populations these proteins are regulated and required, nor how they regulate downstream effectors of epithelial integrity.

The roof plate is a specialized glial cell population that defines the dorsal neural tube/spinal cord midline and patterns dorsal neuroepithelial cells (reviewed in Lee and Jessell, 1999). In mid-embryogenesis, roof plate cells are transformed from wedge-shaped cells into a thin, dense septum of elongated dorsal midline Nestin^(+)^ radial glia (hereafter termed dmNes^+^RG) that extend from the ventricle to the pia (Altman and Bayer 1984; Bohme, 1988; Kondrychyn et al 2013; Sevc et al., 2009; Snow et al., 1990; Sturrock 1981; Xing et al., 2018) and eventually contribute to the EL (Ghazale et al., 2019; Shinozuka et al., 2019; Xing et al., 2018). Elongation of roof plate/dmNes+RG coincides with reduction of the lumen (Bohme 1998; Kondrychyn et al 2013; Korzh, 2014; Sevc et al., 2009; Shinozuka et al., 2019; Xing et al., 2018), and dmNes^+^RG play a critical role in dorsal collapse. Studies in zebrafish suggest that the filamentous (F)-actin cytoskeleton belt that defines the apical side of VL cells, and whose constriction drives early neurulation (Hildebrand and Soriano, 1999; Nagele et al., 1987; Sadler et al., 1982; Sawyer et al., 2010; reviewed in Nikolopoulou et al., 2018) and neuronal delamination (Das and Storey 2014; Kasioulis et al., 2017), is also required for dorsal collapse: this involves a process that depends on its appropriate tethering by elongating roof plate/dmNes^+^RG cells (Kondrychyn et al., 2013; Korzh, 2014). However, in addition to their tethering function, dmNes^+^RG cells may play an active role in promoting delamination by locally regulating VL progenitor cell polarity.

Here we provide evidence that in mouse, Crumbs2 (CRB2) is required for dorsal collapse. In particular, we show that a CRB2-mediated interaction of dmNes^+^RG cells and adjacent VL cells drives the progressive delamination of VL cells and the transformation of the VL to the ependymal layer. Expression of apical/tight junction components and adhesion complexes are maintained at high levels on the apical end-feet of dmNes^+^RG throughout lumen diminution. By contrast, expression of apical/tight junction components and adhesion complexes are dramatically reduced on dorsal-most progenitor cells as they become excluded from the VL. dmNes^+^RG are rich in secretory vesicles and gain-of-function in vivo studies show they secrete a factor that promotes progenitor cell delamination. Further, they express a variant of CRB2 (CRB2S) that can be secreted and appears to mediate this activity. Ex vivo cell culture studies show that CRB2S binds to the apical surfaces of CRB2-expressing epithelial cells, and reduces their cohesion. In vivo analysis of Crb2^F/F^/Nestin-Cre^+/−^ mice confirms an essential role for CRB2 in EL formation. Thus, similar to recent studies on mouse gastrulation (Ramkumar et al., 2016) our data show that CRB2 is required to remove, rather than maintain cells within an epithelium. We propose that collapse is initiated by the release of CRB2S from dmNes^+^RG that acts on CRB2-expressing VL cells, causing downregulation of polarity and junctional proteins and their decreased cohesion. We suggest that these are a first step in exclusion of these cells from the VL and transformation of the VL to the ependymal layer. Our findings suggest a model in which CRB2S acts cell non-autonomously to orchestrate progenitor cell delamination from an epithelium through a mechanism that retains epithelial integrity.

## Results

### Collapse occurs through attrition of dorsal VL cells

Dorsal collapse of the mouse spinal cord occurs over the period E14-E17. At dorsal thoracic levels, the lumen spans almost the entire dorso-ventral length of the spinal cord at E14, reduces to a two-fifth span at E15 and a one-fifth span by E17 (Fig,1A-E; Table S1). As described recently (Cañizares et al, 2019), dorsal collapse is predicted by differences in ventricular layer (VL) morphology. In the ventral VL, the lumen is narrow, nuclei are tightly-packed and medio-laterally oriented (Fig.1A-D white bracket, Supp.Fig.1A-B’), whereas in the dorsal VL, the lumen is wide and nuclei are more loosely-arranged (Fig.1A-C, red bracket; Supp.Fig.1A-B’). The dorsal VL reduces significantly in length on each consecutive day during collapse (Fig.1A-D, red bracket), whereas the ventral VL does not (Fig.1A-D, white bracket; Table S2). Dorsal collapse is the most obvious remodelling event, but is mirrored ventrally by a re-arrangement of floor plate cells, only a subset of which remain within the central canal (Fig.1B, arrowhead; Cañizares et al, 2019).

**Figure 1:**
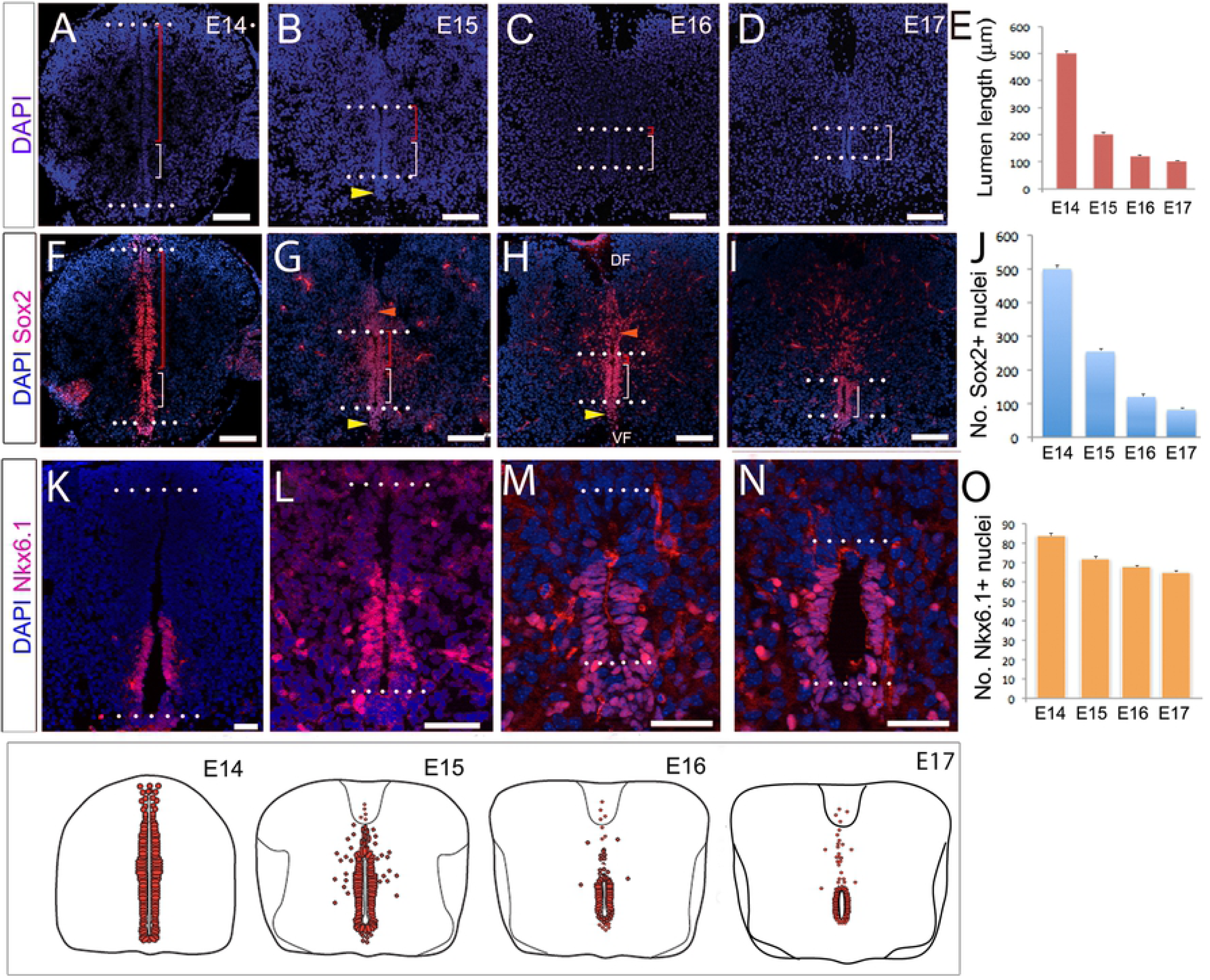
Collapse occurs through attrition of dorsal VL cells. Transverse sections through the spinal cord of E14-E17 mouse embryos. (A-D) DAPI labelling shows diminution of dorsal VL: dotted lines show upper and lower limits of lumen; red bracket indicates dorsal VL; white bracket indicates ventral VL. (E) Quantitative analysis of lumen length. (F-I) Immunolabelling shows Sox2^+ive^ cells throughout the VL, as well as those excluded dorsal to the VL (orange arrowheads) or dissociated ventral to the VL (yellow arrowheads), many in the midline near the ventral funiculus (VF) and dorsal funiculus (DF). (J) Quantitative analysis shows reduction of Sox2^+ive^ VL cells around the lumen. (K-N) Nkx6.1 marks the ventral VL and many Nkx6.1^+ive^ progenitors are retained in collapse (shown quantitatively in (O)). Scale bars: A-D; F-I: 100μm; K-N: 50μm.

SoxB1 genes are expressed on all VL cells during remodelling (Fig.1F-I; Supp.Fig.1C-F; Cañizares et al, 2019). Quantitative analyses show a reduction in the number of Sox2^+ive^ cells in the VL during collapse (Fig.1J; Table S3), and see (Cañizares et al, 2019); excluded cells continue to express Sox1-3 (Fig.1G,H; Supp.Fig.1C-F, orange arrowheads). Likewise, Pax6, which marks cells in all but the ventral-most VL region (consistent with its earlier expression in progenitor subsets; Alaynick et al., 2011), reveals a similar proportional reduction (Supp. Fig.1G-J; Table S3). By contrast, Nkx6.1, which marks ventral progenitor subsets (Alaynick et al., 2011), is restricted to ventral VL cells (Fig.1K), and is then detected at the ventral midline as the floor plate pinches off (Fig.1L-N). There is less proportional reduction in the number of Nkx6.1^+ive^ VL cells (Fig.1; Table S3). Together, these observations are consistent with the idea that dorsal collapse is largely driven through the attrition of the dorsal VL (Fu et al., 2003; Cañizares et al, 2019).

### Dorsal VL cells re-orientate during attrition

The looser arrangement of dorsal VL cells is accompanied by changes in the position and orientation of nuclei that suggest a local cell re-organisation and remodelling of the ventricular layer. Consistently, during dorsal collapse, we detect a nuclear bridge, spanning the two sides of the dorsal VL. Regardless of the stage, the bridge is detected ∼2-8 cell diameters below the dorsal-most lumen (Fig.2A-D, arrowheads). Second, as VL cells dorsal to the bridge are excluded, their nuclei appear to re-orient, from medio-lateral to dorso-ventral (Supp.Fig.1A,A’). By the end of dorsal collapse, nuclei dorsal to the VL are dorso-ventrally elongated (Supp.Fig.1B,B’).

**Figure 2:**
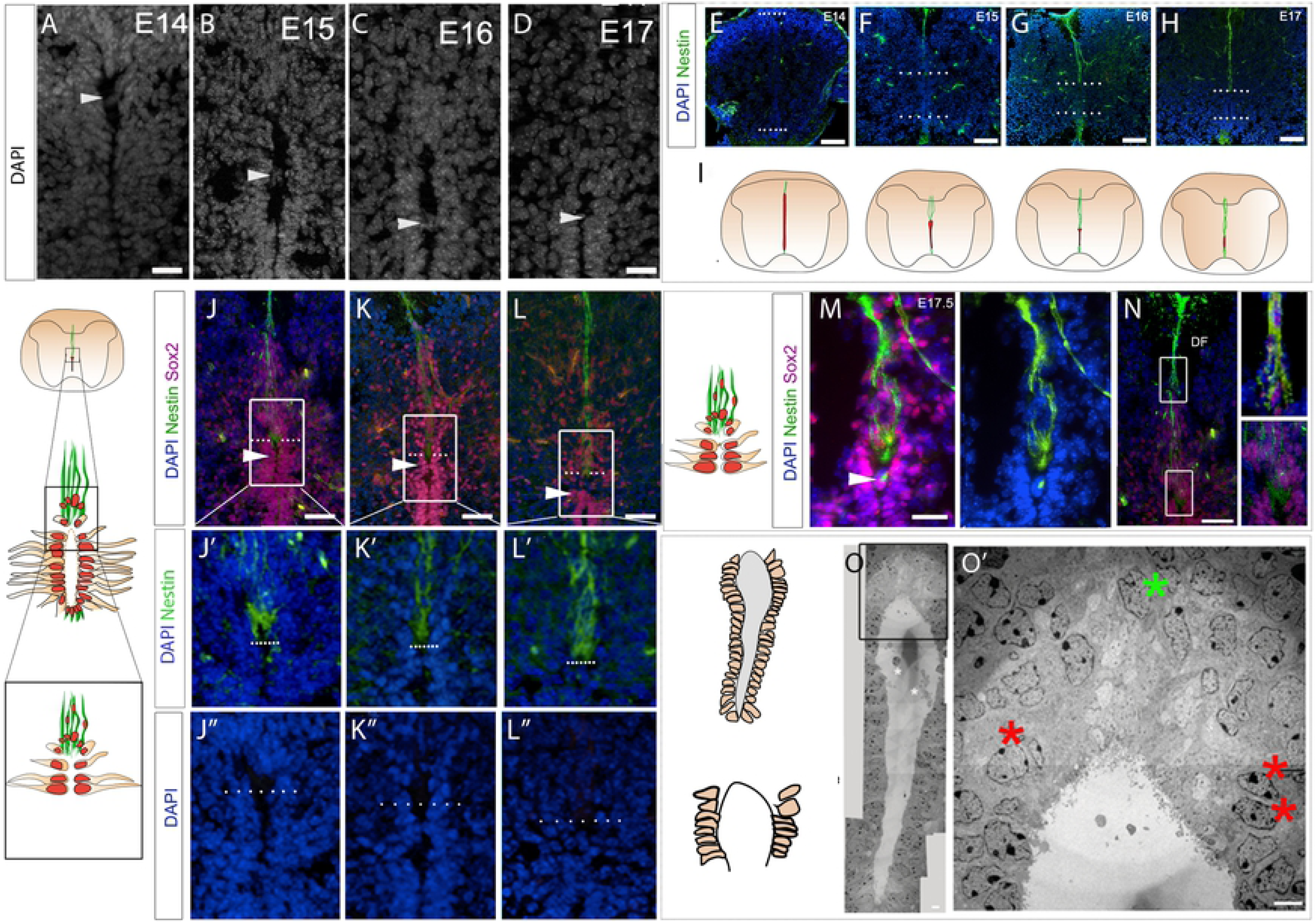
Cell remodelling during dorsal collapse. Transverse sections through the spinal cord of E14-E17 mouse embryos. (A-D) High power views of dorsal VL/lumen; arrowheads point to nuclear bridges. (E-H) Immunolabelling reveals ventral and dmNes^+^RG that elongate as collapse proceeds, shown schematically in (I). (J-L) Co-labelling of Nestin and Sox2 shows that dmNes^+^RG define the dorsal pole. (J’-L’’) High power views (boxed regions in J-L and schematics) reveal that the dorsal-most lumen is occupied by the end-feet of dmNes^+^RG. (M) At E17.5, in addition to expression on dmNes^+^RG, Nestin can be detected in the nuclear bridge below the dorsal lumen (M arrowhead). (N) At E17.5, dorso-ventrally oriented Sox2^+ive^ nuclei are closely apposed to dmNes^+^RG. (O) EM image at E15.5: boxed region shown in O’. Red asterisks show VL nuclei that are close to, but do not abut the lumen; green asterisk shows nuclei that are dorso-ventrally oriented in the dorsal midline some distance above the lumen. Scale bars: A-D: 20μm; A’-D’’: 100μm; B”’-D”’, F: 50μm; B”’’-D”’’, E,E’: 20μm; G: 10μm; G’: 1μm.

To better characterise the position of re-orientating cells, we compared expression of Sox2 to that of Nestin. From E14.5, the dorsal and ventral midline are characterised by Nestin^+ive^ radial glial cells (Fig.2E-I; Supp.Fig.2). Dorsal midline Nestin^+ive^ radial glia (dmNes^+^RG), which elongate as collapse proceeds, are thought to tether the diminishing VL to the pial surface (Snow et al 1990, Sturrock 1981, Bohme, 1988, Altman and Bayer 1984, Sevc 2009, Kondrychyn et al 2013; Cañizares et al, 2019; Xing et al., 2018; Shinozuka et al., 2019). Therefore Nestin expression marks the dorsal pole of the diminishing VL.

Double-labelling of Nestin and Sox2 confirms that the dorsal-most lumen is occupied largely by dmNes^+^RG end-feet: no nuclei could be detected in this region (Fig.2J-L”; Snow et al., 1990). Further, it confirms that Sox2^+ive^ cell bridges are found ∼2-8 cell diameters below dmNes^+^RG end-feet, ie ∼2-8 cell diameters below the dorsal lumen (Fig.2J-L). Towards the end of dorsal collapse (E17.5), a bright spot of Nestin immunoreactivity is detected in the Sox2^+ive^ bridge (Fig.2M,M’), suggesting a physical interaction of dmNes^+^RG and dorsal VL cells. Double-labelling of Nestin and Sox2 confirms, additionally, that many Sox2^+ive^ cells are closely associated with dmNes^+^RG and can be detected as far away as the dorsal funiculus (Fig.2N).

To validate the arrangement/re-organisation of dorsal VL cells, we performed transmission electron microscopy (EM) imaging at E15.5-E16 (n=3 embryos; 6 sections), a time when dorsal collapse is underway. The narrower ventral VL and wider dorsal VL zones can be distinguished in EM images (Fig. 2O). Quantitative analyses confirmed the looser arrangement of dorsal VL cells (21.5+/-1.04 nuclei/100μm^2^) to ventral VL cells (54.0+/-1.83 nuclei/100μm^2^ respectively; Table S4: significantly different; p<0.0001), and high power views confirm that the dorsal-most pole lacks nuclei (Fig.2O’).

Notably, however, although nuclei in the dorsal VL are loosely arranged, they remain abutted to the lumen except for the 2-3 nuclei directly adjacent to the dorsal pole. These appear to have moved away (Fig 2O’; red asterisks), suggesting an organised delamination at the dorsal end.

In summary, a first indication of dorsal collapse is a local cell re-organisation and remodelling of the dorsal ventricular layer. Key changes include formation of a bridge just below the dorsal-most pole, the re-orientation of nuclei dorsal to the bridge, and distancing of nuclei from the lumen in cells that are immediately proximal to dmNes^+^RG.

### Dorsal collapse involves cell delamination and ratcheting

To better examine the rearrangement of cells during dorsal collapse we performed timelapse imaging of spinal cord slice cultures, focusing on the dorsal regions of the collapsing spinal cord. After transfecting sparse numbers of cells with membrane-GFP and histone-RFP (n=3 slices from 3 embryos; 6-12 dorsal cells electroporated in each), the dmNes^+^RG cell can be detected due to its typical morphology, midline position, and basal nucleus (detected in all 3 slices; Fig.3A yellow arrowhead; Frame 0). As predicted from our static studies (Fig.2), dmNes^+^RG cells remain in a dorsal midline position throughout imaging (Fig.3A Timeframes 0 to 1060; Movie 1; 6 cells observed from 3 slices). At their apical ends, dmNes^+^RG cells are closely-apposed to a dorsal VL progenitor cell (Fig.3A Timeframe 0 white arrowhead). This cell undergoes apical abscission (Fig.3A asterisks Timeframe 440; Das and Storey, 2014); the apically-retained fragment is initially large, but gradually reduces/disappears (Fig.3A yellow asterisks Timeframe 440 to 700). A second VL progenitor cell then becomes closely-apposed to the dmNes^+^RG cell (Fig.3A Timeframe 990 arrowhead; total of 9 step-wise abscising cells observed from 3 slices). Together this suggests that dorsal collapse proceeds through the progressive abscission of dorsal VL progenitors that are immediately adjacent to the dmNes^+^RG cell, and the ratcheting down of the Nestin^+ive^ end-foot, to the next VL progenitor. This progressive series of abscission events and accompanying ratcheting down of the dmNes^+^RG cell population suggests a novel and specific delamination mechanism.

**Figure 3:**
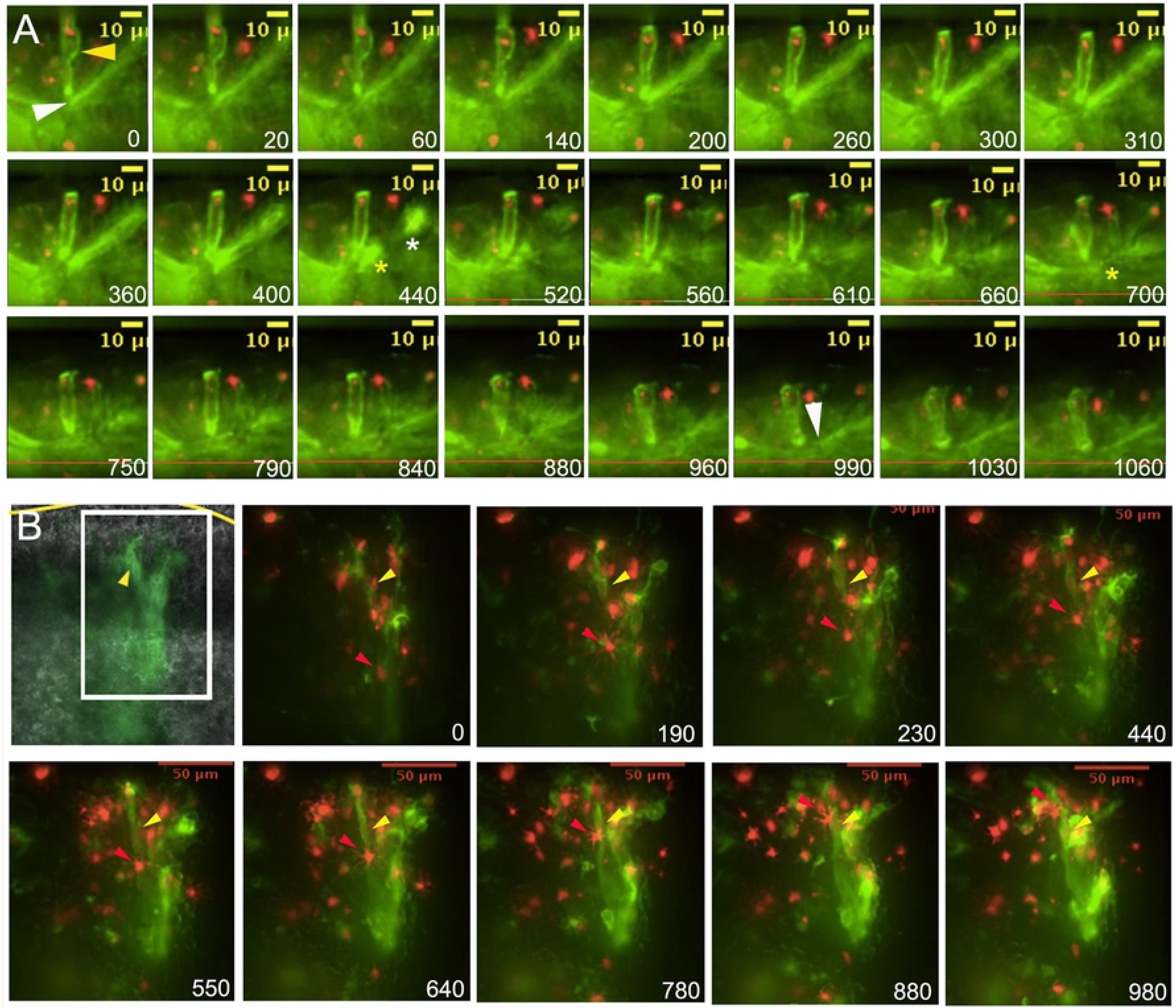
Dorsal collapse involves cell delamination and ratcheting. (A) Sequential stills from time-lapse imaging after low density electoporation of membrane-GFP histone-RFP. A midline cell, whose morphology indicates it to be a dmNes^+^RG remains throughout the culture (yellow arrowhead, Timeframe 0). An adjacent cell, apically attached (white arrowhead, Timeframe 0) undergoes abscission (white asterisk shows main body of delaminating cell; yellow asterisk shows apical remnant, Timeframe 440). The apical remnant is lost (yellow asterisk, Timeframe 700) and a second VL cell comes to abut the dmNes^+^RG (white arrowhead, Timeframe 990). (Movie 1). (B) Sequential stills from time-lapse imaging after high density electoporation of membrane-GFP histone-RFP. Yellow arrowhead points to a dmNes^+^RG, whose position remains the same throughout the culture. Red arrowhead points to a nucleus that migrates dorsally. (Movie 2).

Post-abscission, labelled cells appear to migrate dorsally. To examine this further, larger numbers of cells were transfected with membrane-GFP and histone-GFP (n= 2 slices; 20-30 dorsal cells electroporated in each). Although, in these cases, the apical parts of cells could not be resolved, many dmNes^+^RG cells could be detected in the dorsal midline (Fig.3B). Associated with these were nuclei that moved dorsally, apparently using the processes as a scaffold for dorsal migration (Fig.3B, red arrowheads; Movie 2. This suggests that after their abscission, VL progenitors actively migrate dorsally along the dmNes^+^RG scaffold, rather than remaining localised on the elongating dmNes^+^RG cell.

### Diminished adhesion junctions on VL progenitors adjacent to dmNes^+^RG

The stable retention of the dmNes^+^RG at the dorsal pole and abscission of immediately-adjacent VL progenitors led us to predict that these two cell types will show distinctive profiles of apical adhesion and junction proteins that, in other systems, govern epithelial integrity and cell delamination (reviewed in Pichaud, 2018). We therefore examined expression of the apical polarity proteins aPKC, Crumbs2 (CRB2) and PAR3, and Zona occludens 1 (Zo-1), a tight junction-associated protein, at E13.5 (prior to collapse), E15-E15.5 (maximal dorsal collapse and E17 (termination of collapse). At the same time we assayed expression of phalloidin, a marker of F-actin, previously suggested to anchor dmNes^+^RG and VL progenitor cells (Kondrychyn et al., 2013). At E13.5, junction/adhesion proteins are detected in a continuum on the apical side of all VL progenitor cells (Fig. S3A-C). By contrast, at E15, junction/adhesion proteins are discontinuous on dorsal VL cells. High levels of expression of Zo-1, aPKC, CRB2 and PAR3 are detected on the end-feet of dmNes^+^RG (Fig.4A-D, A’-D’, A”-D” open arrowheads). However, Zo-1, aPKC, PAR3 and CRB2 are not expressed, or are expressed in a discontinuous manner on dorsal VL cells (Fig.4A-D, A’-D’, A”-D”). Most notably, Zo-1, aPKC and PAR3 cannot be detected on dorsal VL progenitor cells that lie immediately adjacent to the dmNes^+^RG (Fig.4A-A”,B-B” D-D” green arrowheads; n=24-30 cells each, imaged from minimum of 3 embryos Supp. Fig.3D), while CRB2 appears absent/diminished or reduced to a single dot on these cells (Fig.4C-C”; n=24 cells from 8 embryos; Supp. Fig.3). By E17, some dorsal discontinuity is still apparent, albeit less obvious (Fig.4KF-I; Fig.S3E; n=8 slices from 4 embryos), and by E18, apical/junctional proteins are again expressed as a continuum (Fig.S3F). Phalloidin does not show the same marked absence, but shows punctate labelling on the apical side of dorsal VL cells at E15 (Fig.4E-E”; Supp. Fig.3D), and then continuous labelling at E17 (Fig. 4J). Together these data suggest that apical adhesion/tight junction proteins are retained on dmNes^+^RG throughout collapse, but are reduced on dorsal VL progenitor cells; in particular, they are absent (Zo-1, aPKC, PAR3) or markedly reduced (CRB2) from the dorsal cells that are delaminating.

**Figure 4:**
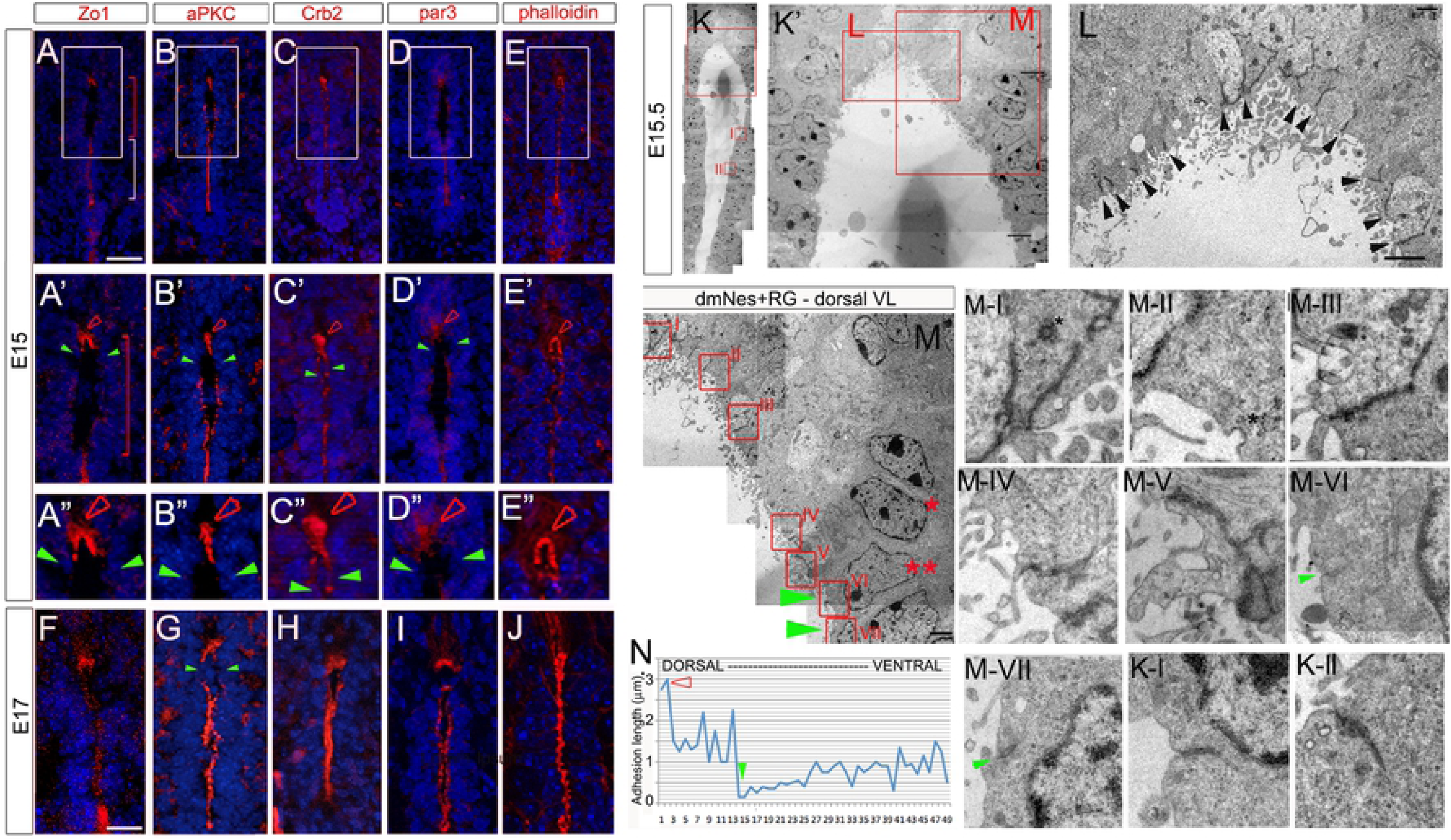
Apical polarity proteins and tight junctions are reduced on delaminating VL cells. (A-E”) Transverse sections through spinal cord of E15 mouse embryos immunolabelled as indicated. (A’-E’) show high power views of boxed regions shown in (A-E); (A”-E”) show high power views of regions indicated by arrowheads in (A’-E’). High expression of apical polarity proteins and the tight junction protein, Zo-1, are detected on the end-feet of dmNes^+^RG (open red arrowheads), but not detected (Zo-1, aPKC, PAR3) or barely detected (CRB2) on immediately adjacent dorsal VL cells. Phalloidin is expressed strongly on dmNes^+^RG, and in a punctate manner throughout dorsal VL cells. (F-J) At E17, junctional and polarity proteins likewise show reduced/no expression on VL cells that abut dmNes^+^RG but phalloidin is detected in a continuum. (K-M) EM images of the E15.5 VL. Boxed regions in (K) shown in (K’, K-I and K-II). Boxed regions in (K’) shown in L, M). In (L) arrowheads point to electron-dense junctions on dmNes^+^RG. In (M), red asterisks point to nucleus about to delaminate (double asterisk) or just delaminated (single asterisk); red boxed regions in M are shown at high power in panels M-1 to M-VII. Electron-dense junctions are barely detected in cells that are about to delaminate (M-VI and M-VII; green arrowheads point to junctions). Asterisks in M-I and M-II point to secretory vesicles. (N) Quantitative analysis showing length of electron-dense junctions along a single side of the VL: junction length increases in a gradient, dorsally and ventrally, from delaminating cells. Scale bars: A-J:100μm; K-M: 2μm.

To explore this further, we examined adhesion complexes after EM imaging (Fig.4K,K’). High power views of dorsal-most regions show that dmNes^+^RG endfeet have long, tight adhesion complexes, whose characteristic profiles (electron-dense; extending from the apical surface on adjacent cell membranes) can be detected at high-power (Fig.4L arrowheads, 4M boxed regions M-I to M-III). By contrast, adhesion complexes can barely be detected on VL progenitor cells that are about to undergo abscission and delamination: only electron-lucent, short adhesion complexes can be detected (Fig 4M boxed regions M-VI and M-VII, green arrowheads). Immediately dorsal to these, a long adhesion complex extends parallel to the VL (Fig.4M-V). In VL progenitor cells that lie ventral to the abscising cell, adhesion complexes lengthen and become increasingly electron-dense (Fig.4 K boxed regions K-I, K-II; Fig.4N).

In summary, throughout the collapse window, dmNes^+^RG form long apical adhesion complexes and tight junctions. By contrast VL progenitors that are about to abscise show markedly reduced apical adhesion complexes and tight junctions. Dorsal and ventral to these, apical adhesion complexes gradually increase in length. Together these analyses show that a reduction in apical adhesion/tight junction complexes prefigures or accompanies VL progenitor delamination.

### dmNes^+^RG secrete a factor that promotes VL progenitor delamination

dmNes^+^RG have previously been implicated in dorsal collapse: studies in zebrafish have suggested that dmNes^+^RG cells are tethered to VL cells via a filamentous (F)-actin cytoskeleton belt, and that tethering is required for dorsal collapse (Kondrychyn et al., 2013). However, our EM imaging shows that dmNes+RG are rich in vesicles that appear to be fusing with the adjacent lumen (Fig.4M-I, 4M-II asterisks; Shinozuka et al., 2019), suggesting that these cells could be secreting a factor involved in VL progenitor delamination.

To test this idea, we established an in vivo assay, using the early chicken embryonic neural tube to assay dmNes^+^RG activity. Like mice, chicken embryos undergo collapse (between E8 and E11; Whalley et al., 2009; Wakamatsu 2007) and have dmNes+RG cells (Supp.Fig.4), suggesting they have a conserved cellular machinery that will allow them to respond to mouse dmNes^+^RG cells. Mouse E15 dmNes^+^RG cells, or control VL cells, were transplanted into the dorsal lumen of HH st10 (E1.5) chick embryos, a stage when the neural tube is composed of pseudostratified neuroepithelial cells, and embryos were developed 20-24 hrs, to E2.5 (Fig.5A schematic; n=14 embryos each). In embryos transplanted with lateral VL cells, the neural tube appeared normal: laminin or dystroglycan-labelling showed that the basement membrane was intact and the distribution of Pax6^+ive^ and Nkx6.1^+ive^ progenitors appeared normal (Fig.5C,D). Shh was detected on the floor plate and notochord (Fig.5E). Zo-1 and aPKC were both detected in an unbroken manner on apical VL progenitor cells (Fig.5F,G) (Table S5). By contrast, in chick embryos transplanted with dmNes^+^RG, neuroepithelial cells were disrupted: dystroglycan- and laminin- labelling showed breaks in the basement membrane (Fig.5H, I). Within the neural tube, Pax6^+ive^ and Nkx6.1^+ive^ progenitor cells were more broadly distributed than in controls, and appeared disorganised (Fig.5H,I); some Pax6^+ive^ cells appeared to be delaminating from the neuro-epithelium and were detected outside of the basement membrane boundary (Fig.5H, arrowheads; Table S5). Ectopic clumps of Shh^+ive^ cells were similarly detected outside of the neural tube: these appeared close to the endogenous floor plate (Fig.5J, arrowheads) and in some cases, appeared to be dissociating from it (Supp.Fig.5A,B; Table S5). In embryos analysed 20h post-operatively, Zo-1 and aPKC were downregulated from the apical side of neuroepithelial cells (Fig.5K,L). To further test the specificity of this effect, we examined a wider range of tissues. Nestin^+ive^ cells in dorsal regions of the subventricular layer (SVL) of the lateral ventricle evoked a similar response to dmNes+RG cells (Supp.Fig.5C-G; Table S5). By contrast, other regions of the CNS, including ventral RG failed to mimic E15 dmNes^+^RG tissue (Table S5).

**Figure 5:**
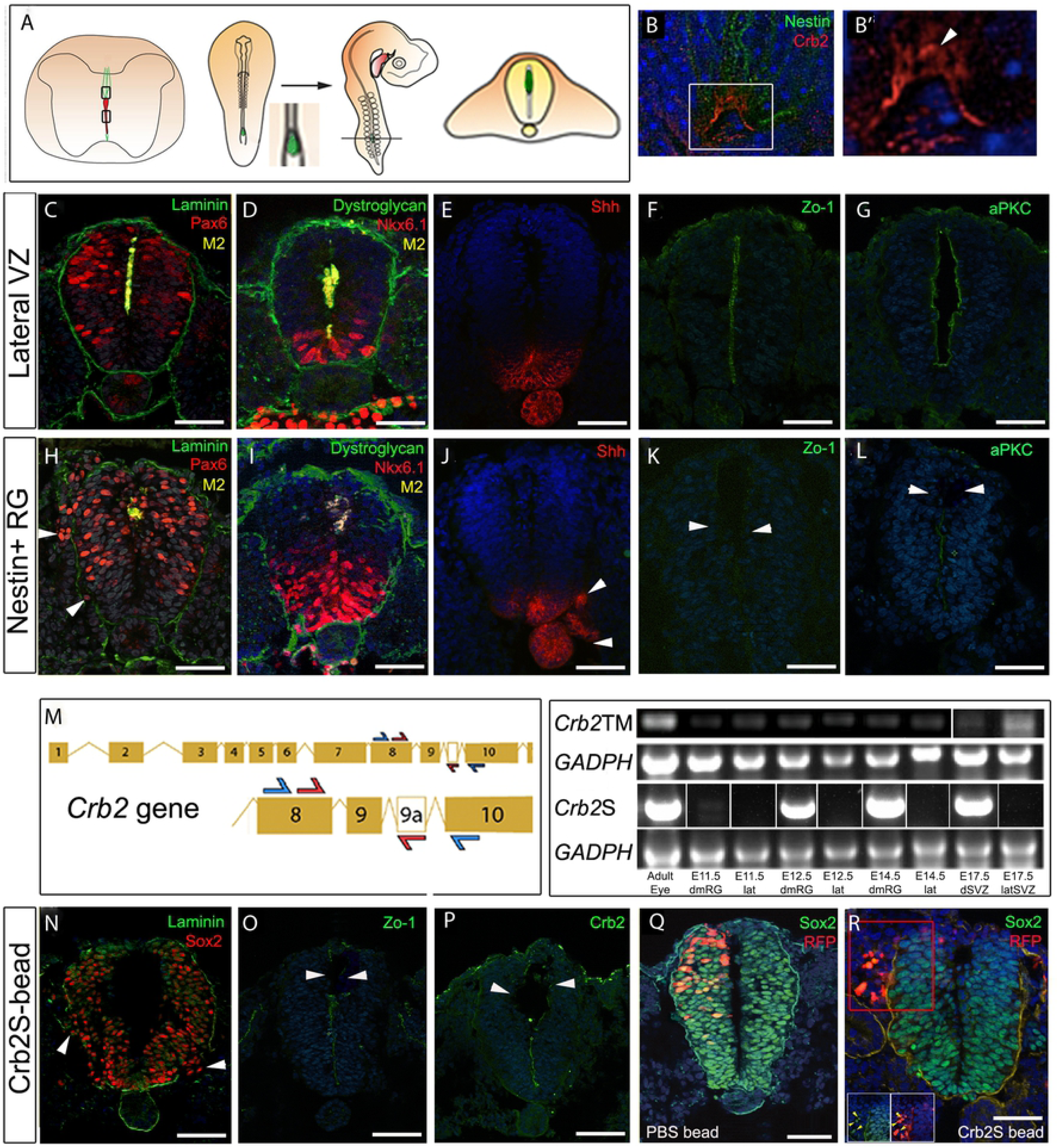
CRB2S from dmNes^+^RG promotes delamination. (A)Schematic showing the grafting procedure: dmNes^+^RG or VL cells dissected and grafted to HH st10 chick embryos. After 20-24h incubation (to HH st16-18), chicks sectioned in operated region. (B-B’) Transverse sections of E16 mouse spinal cord, double labeled to detect Nestin and CRB2. (B) shows confocal view: single channel view shown in (B’); arrowhead in B’ points to non-apical CRB2 expression. (C-G) Control tissue grafts, immunolabelled as shown. (C,D) Serial adjacent sections after a 24h incubation. Mouse tissue is detected in the lumen; Pax6 and Nkx6.1 are detected in the neural tube; the basement membrane is intact, revealed through laminin or dystroglycan labelling (E-G) Serial adjacent sections after a 20h incubation. Shh is detected on notochord and floor plate; Zo-1 and aPKC are detected on the apical side of neuroepithelial progenitor cells. (H-L) dmNes^+^RG grafts. (H,I) Serial adjacent sections after 24hr incubation: basement membrane is disrupted, Pax6^+ive^ neural progenitors are detected ectopically outside the disrupted basement membrane (H, arrowheads) and Nkx6.1^+ive^ progenitors are disorganised. (J-L) Serial adjacent sections after a 20h incubation. Ectopic Shh^+ive^ cells are detected outside of the neural tube (J, arrowheads); little/no expression of Zo-1 and aPKC is detected (K,L). (M) Left hand panel: Schematic showing primer strategy against *Crb2* or *Crb2S*. CRB2S has an additional exon alternatively spliced into the transcript. Blue arrows indicate a first round of PCR, amplifying a band between exon 8 and 10 present in both *Crb2* cDNA and *Crb2S* cDNA. Nested primers are indicated by red primers, which amplify between exon 8 and exon 9a, present in only *Crb2S*. Right hand panel: Amplification using first round and nested primers, of adult eye, E11.5, E12.5 and E14.5 dmNes^+^RG (dmRG), E11.5, E12.5 and E14.5 lateral VL (lat), and dorsal or lateral E17.5 SVZ samples. A GADPH control is provided for both sets of primers in all samples. *Crb2S* lanes show representative images from 4 biological samples: see Fig.S7 for full gel. (N-P) Transplanted CRB2S protein-soaked bead grafted to HH st10 chick embryo analysed after 24hrs. The basement membrane is disrupted and Sox2^+ive^ neural progenitors are detected outside the disrupted basement membrane (N, arrowheads). Zo-1 and CRB2 expression are reduced in the vicinity of the bead (O, P, arrrowheads). (Q) Control RFP electroporated and PBS-soaked bead-grafted HH st10 chick embryo after 24hrs shows normal expression of Sox2 throughout neural tube and normal basement membrane. (R) RFP electroporated and CRB2S-soaked bead grafted HH st10 chick embryo after 24hrs shows ectopic expression of Sox2 outside neural tube and disrupted basement membrane. Red box indicates inset below. Yellow arrows indicate Sox2^+ive^ RFP^+ive^ cells outside neural tube. Scale bars: C-L, N-R: 50μm; B-B’: 10μm.

The ability of transplanted dmNes+RG (and SVL cells) to affect progenitor cells at a distance indicated that these cells may secrete a factor that leads to downregulation of apical polarity and tight junction proteins. Further, this factor appears to initiate events that lead to the delamination of neuroepithelial cells.

### *Crb2S* is expressed in dmNes^+^RG and CRB2S mimics their activity

We had noted that during the period of dorsal collapse, CRB2 appears more diffuse in the apical endfeet of dmNes^+^RG than other apical/junctional proteins (Fig.5B, B’) and that similarly punctate expression of CRB2 is detected in the Nestin^+ive^ SVL cells of the lateral ventricle (Supp. Fig.5B,B’). Studies in humans suggest the existence of an alternatively spliced isoform of CRB2, that lacks the transmembrane domain and is putatively secreted (Katoh and Katoh, 2004). This prompted us to investigate whether a secreted isoform of CRB2 can be detected in mouse, and whether this protein can phenocopy the effects of dmNes^+^RG cell transplantation.

Bioinformatic analysis of the mouse *Crb2* locus predicts different splice variants, in one of which, alternative exon splicing between exon9 and exon10 introduces a premature stop codon before the transmembrane domain, predicting that this splice variant (which we term *Crb2+9A*) may encode the putatively secreted protein. Stable clonal HEK293 cell lines constitutively expressing *Crb2+9A* secreted a CRB2-protein (termed CRB2S) into the media (Supp.Fig.6 and Methods). We therefore used a nested PCR approach (Fig.5M) to determine whether the mRNA encoding the secreted isoform can be specifically detected in dmNes^+^RG cells. dmNes^+^RG, SVL cells, and lateral VL cells were compared to the adult eye, a tissue that expresses high levels of transmembrane CRB2 (van de Pavert et al., 2004; Paniagua et al., 2015). As predicted from immunohistochemical analyses, RNA encoding full-length transmembrane CRB2 was detected in both dmNes^+^RG cells and lateral VL cells. However, the RNA encoding the secreted isoform, CRB2S, was detected only in dmNes^+^RG and not in lateral VL cells (Fig.5M; Supp.Fig.7).

Beads, soaked in purified CRB2S, were then implanted into the dorsal lumen of HH st10 chick embryos (n=6). Transplants of beads soaked in CRB2S phenocopied transplants of dmNes^+^RG cells: disruption and disorganisation of the neuroepithelium; breaks in the basement membrane; reduction of Zo-1, CRB2 and aPKC, most obviously in the vicinity of the bead (Fig.5N-P arrowheads); the ectopic appearance of Sox2+^ive^, Pax6^+ive^, Nkx6.1^+ive^ and Shh^+ive^ neuroepithelial cells outside of the neural tube (Fig.5N; Supp.Fig.8 arrowheads; Table S5). PBS soaked beads caused no disruption to the chick neuroepithelium (Supp.Fig.8; Table S5).

The appearance of ectopic progenitor and floor plate cells outside of the neural tube could arise, either through the delamination of neuroepithelial cells, or due to a fate change in cells outside of the neural tube. To distinguish between these, we combined RFP electroporation and bead implantation. When RFP was electroporated into the dorsal neural tube (avoiding the neural crest) prior to transplantation of a PBS bead, Sox2^+ive^ RFP^+ive^ cells were confined to the neural tube (Fig.5Q). By contrast, when similar cells were targeted, then a CRB2S bead implanted, Sox2^+ive^ RFP^+ive^ cells were detected outside of the neural tube (Fig.5R). Thus, in the presence of CRB2S, cells delaminate from the neuroepithelium. In summary, a secreted variant of CRB2, CRB2S, is specifically expressed in dmNes^+^RG cells, and its premature mislocalisation leads to loss of apical polarity and junctional complexes in neuroepithelial cells, and their ability to delaminate from the neuroepithelium.

### CRB2S reduces cohesion/adhesion in MDCK cells

CRB2 can bind to itself in a homophilic interaction in epithelial cells, and disruption of CRB2-CRB2 leads to a loss of polarity and disrupted epithelial-mesenchymal transition (EMT) (Zou et al., 2012). This raises the possibility that CRB2S can similarly associate with CRB2. To begin to test this, we incubated spinal cord sections with His-tagged CRB2S, then analysed with an anti-His antibody. Punctate labeling was detected on the apical side of VL cells (Fig.6A).

**Figure 6:**
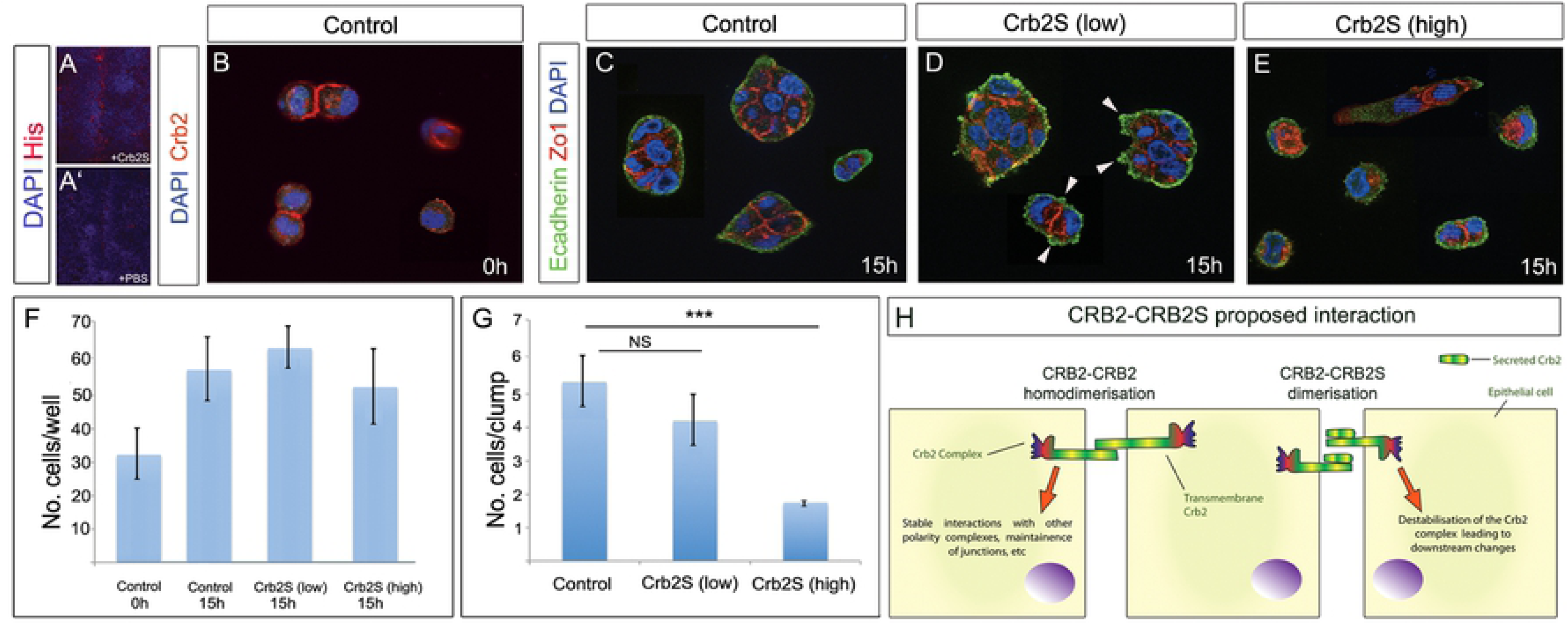
CRB2S reduces cohesion/adhesion in MDCK cells. (A,A’) Transverse sections of E15.5 spinal cord, incubated with His-tagged CRB2S (A) or PBS (A’). Anti-His antibody detects CRB2S at the apical side of VL cells. (B) MDCK cells seeded at low density express CRB2. Expression is detected at junctions between doublets. (C-E) MDCK cells seeded at low density, then cultured an additional 15h in control medium (C), low concentration of CRB2S (D) or high concentration of CRB2S (E). (F,G) Quantitative analyses shows that CRB2S does not affect the total cell number, but significantly reduces the number of cells/clumps. (H) Model for effect of CRB2S. In the absence of CRB2S, transmembrane CRB2 binds homophilically (Zou et al, 2011) to promote stable interactions with components of the apical polarity/junctional complex and maintain neuroepithelial junctions. CRB2S competes homophilically to bind to CRB2. CRB2-CRB2S dimerisation destablilises the CRB2 complex, leading to downstream neuroepithelial destabilization and delamination.

To test the effect of CRB2S on epithelial cells, we performed an acute assay on MDCK cells (a columnar epithelial cell line (Cohen et al., 2004)), dissociated and seeded at low density. DAPI-labelling revealed the presence of single cells and doublets (Fig.6B), and immunohistochemical analysis confirmed that MDCK cells express CRB2 (Fig.6B). Exposure of cells to control medium or to CRB2S for 15h suggested that CRB2S did not affect proliferation or apoptosis of MDCK cells: the total numbers of cells in each condition was not significantly different, and in each condition, a similar increase in total cell number was detected over the 15h culture period (Fig.6F). However, CRB2S had a negative effect on cohesion/adhesion. In control medium, the majority of cells were detected in small clumps (Fig.6B,G). Apically, cells expressed the tight junction protein, Zo-1, and basally, they expressed E-cadherin in a uniform manner (Fig.6C). After exposure to low concentrations of CRB2S, similar-size clumps of cells were detected (Fig.6D,G). However, E-cadherin labelling revealed blebs, associated with unusual cytoplasmic Zo-1 expression (Fig.6D, arrowheads). More strikingly, after exposure to higher concentrations of CRB2S, cells were largely detected as singlets or doublets, with almost no larger clumps detected (Fig.6E,G). As with low CRB2S, Zo-1 showed unusual cytoplasmic expression, and E-cadherin was no longer uniformly detected in cells (Fig.6E). These observations suggest that CRB2S can reduce cohesion/adhesion in CRB2-expressing cells. Together, these analyses suggest a direct interaction of CRB2S and CRB2 and a model in which CRB2S can destabilise CRB2-CRB2 interactions, leading to downstream changes, including loss of polarity and decreased cohesion/adhesion (Fig.6H, schematic: see Discussion).

### *Crb2* is required for dorsal collapse

This model predicts that CRB2 is essential for dorsal collapse. To test this we deleted *Crb2* from neuroepithelial VL progenitors by crossing a transgenic *Crb2* floxed (*Crb2^F/F^*) mouse (Alves et al., 2013) with *Nestin-Cre^+/-^* mice (Dubois et al., 2006) to obtain *Nestin-Cre^+/-^/CRB2^F/F^*) animals (termed CRB2 Nestin-Cre hereafter). Embryos were compared to *Crb2^F/F^* embryos at E17 (n=4 embryos each). In control (*Crb2^F/F^*) embryos, collapse occurred as normal. DAPI labelling showed the lumen was reduced to a similar extent to that seen in wild type mice (compare Fig.7A and Fig.1A) and dmNes^+^RG cells elongated to a similar extent to that in wild-type embryos (compare Fig.7B and Fig.S2); CRB2 itself, aPKC and PAR3 were detected on the apical side of VL cells (Fig.7C-E) and Pax6^+ive^ and Nkx6.1^+ive^ progenitors were located as in wild type embryos (Fig.7F,G).

**Figure 7.**
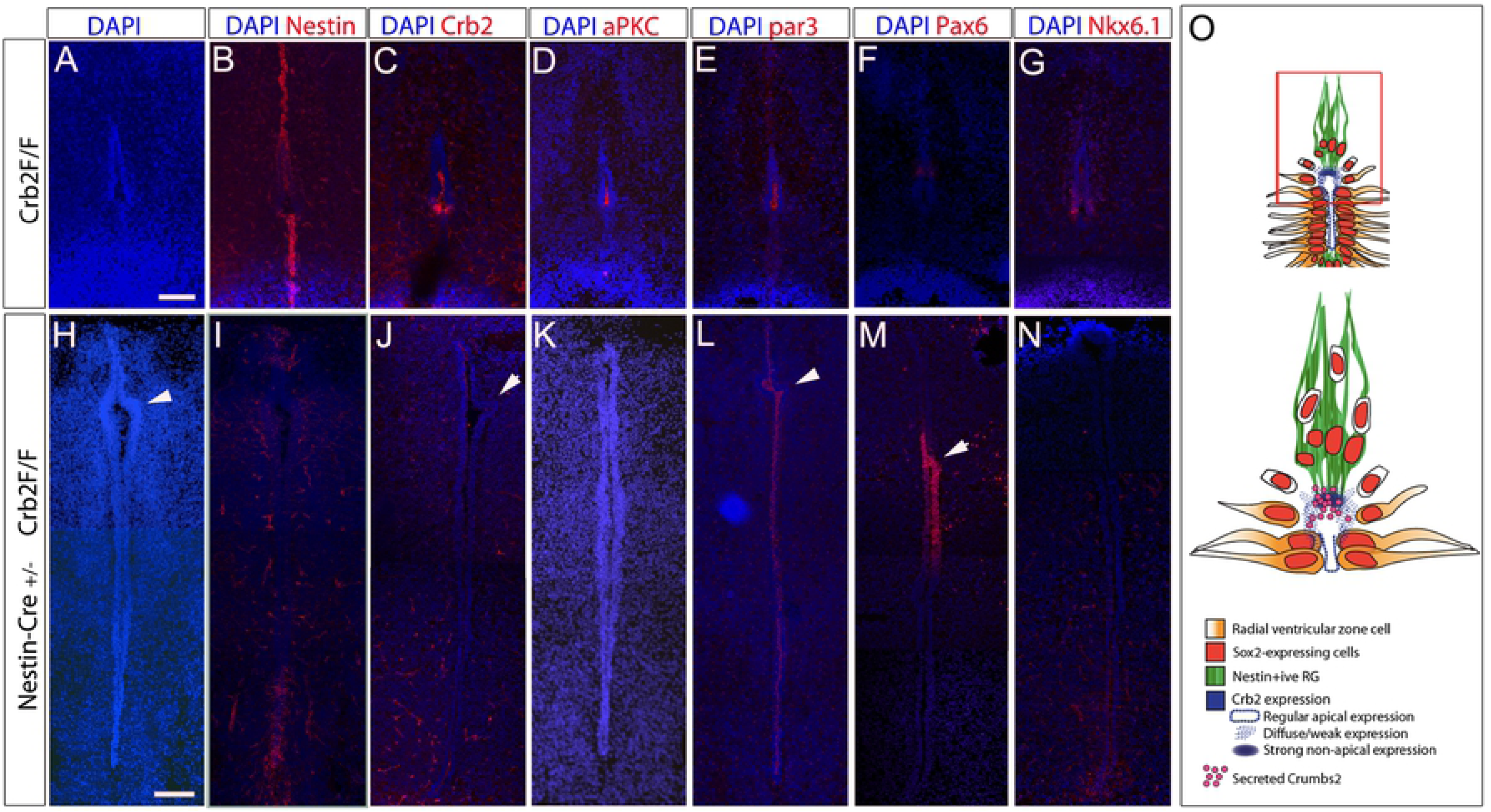
*CRB2* is required for dorsal collapse. Transverse sections through E17 spinal cord in *CRB2^F/F^* control embryos (A-G) or *Nestin-Cre+/- CRB2^F/F^* embryos (H-N), after immunolabelling. Knock-out of transmembrane CRB2 prevents collapse : a long lumen is detected, with unusual kinks (arrowheads). (O) Schematic shows model: CRB2S is secreted from the endfeet of dmNes^+^RG, and binds to adjacent apical cells, resulting in their downregulation of apical/junctional proteins and delamination. Scale bars: 50μm.

By contrast, collapse failed to proceed normally in CRB2 Nestin-Cre embryos. DAPI labelling showed the DV extent of the lumen to be much greater, consistent with a dorsal collapse failure (Fig.7H), although unusual kinks were detected in dorsal regions (Fig.7H,J,L,M arrowheads). Nestin immunolabelling showed that the ventral and dorsal RG failed to elongate (Fig.7I). Analysis with anti-CRB2 Ab confirmed the lack of expression of CRB2 at the apical surface of VL cells (Fig.7J). No expression of aPKC was detected (Fig.7K) but PAR3 was expressed at the apical side of VL cells (Fig.7L). Pax6^+ive^ progenitors were retained in dorsal parts of the VL (Fig.7M), and in contrast to wild-type and *CRB2^F/F^* embryos, no Nkx6.1 expression could be detected on ventral VL progenitors (Fig.7N). Together, this data suggests that CRB2 is required for collapse of the embryonic VL and the establishment of the adult-like ependymal zone.

## Discussion

Here we present evidence for a novel role for the apical protein CRB2 in a local delamination and ratcheting mechanism that remodels the VL of the vertebrate spinal cord. CRB2 is required to effect progenitor cell delamination during dorsal collapse, a process that contributes to central canal formation through attrition of VL cells without loss of overall VL integrity. Mouse embryos that lack CRB2 in Nestin^+ive^ radial glial cells form a neural tube, but VL progenitors fail to delaminate, the lumen does not collapse and the central canal fails to form. Previous studies have revealed that mouse CRB2 is required for gastrulation (Xiao et al., 2011; Ramkumar et al., 2015), and similarly promotes cell ingression during an EMT that generates the germ layers (Ramkumar et al., 2016). However, in this ingression mechanism, classic markers of apico-basal polarity remain in their correct pattern (Ramkumar et al., 2016). In contrast, we find that mouse embryos that lack CRB2 show aberrant apico-basal polarity. Thus, as in the best-defined role of *Drosophila* Crumbs protein (reviewed in Pichaud et al., 2018) and the mouse telencephalon (Dudok et al., 2016), CRB2 acts a determinant of polarity in the neuroepithelium. However, unlike the classic action of Crumbs in *Drosophila*, CRB2 plays an essential role in removing, rather than maintaining, apical junctions. Our studies point to a cell non-autonomous role for a secreted variant, CRB2S, in this mechanism.

CRB2 is detected in a dynamic pattern in cells that line the spinal cord lumen. dmNes^+^RG that tether the lumen to the pial surface maintain high levels of CRB2 in their apical endfeet throughout collapse. By contrast, although CRB2 is detected on the apical sides of all VL cells prior to, and following collapse, it is eliminated from the apical side of VL cells that immediately neighbour dmNes^+^RG during collapse. Full-length transmembrane *Crb2* is expressed in all VL cells, but a splice variant that lacks the transmembrane domain and is predicted to be secreted (Katoh and Katoh, 2004; van der Hurk et al 2005; Watanabe et al., 2004) is specifically expressed in dmNes^+^RG. Three lines of evidence suggest that CRB2S, secreted from dmNes^+^RG, acts on immediately adjacent VL cells to disrupt apico-basal polarity and effect their delamination from the VL. First the CRB2 splice variant can be secreted, and premature exposure of chick neuroepithelial VL cells to CRB2S leads them to lose apical aPKC - mimicking the phenotype of the Crb2 Nestin-Cre cKO mice - and to delaminate. Second, exposure of CRB2-transfected MDCK cells to CRB2S disrupts the basal protein, E-cadherin and the tight junction protein, Zo-1 (similarly reduced in chick neuroepithelial cells after exposure to CRB2S) and reduces their cohesion/adhesion. Finally, the phenotype elicited by CRB2S is recapitulated by dmNes^+^RG: premature and ectopic exposure of chick neuroepithelial VL cells to dmNes^+^RG, but not other spinal cord VL cells, leads to the loss of Zo-1, aPKC, disruption of the basement membrane and delamination of progenitor cells. It remains to be proven that CRB2S is secreted in vivo, but these observations, together with the secretory nature of dmNes^+^RG and their specific expression of the *Crb2* splice variant, provide strong evidence for this idea.

We demonstrate that CRB2S associates with CRB2 on the apical side of VL cells and propose that this interaction disrupts a molecular pathway that maintains neuroepithelial integrity. In flies and vertebrates, full-length CRB2 homologues normally mediate cell adhesion through homophilic interactions at opposing cell membranes, mediated by the extracellular domain. Homophilic interactions stabilise Crb/CRB2 apically, and maintain epithelial organisation and integrity (Omori et al., 2006; Hsu et al., 2010; Zou et al., 2012; Roper, 2012; Letizia et al., 2013; Pichaud, 2018; Richard et al., 2006; Alves et al., 2014). The full mechanisms through which this is achieved remain elusive, but likely involve the ability of CRB2 to interact with the PAR complex (Cdc42-Par6-aPKC-PS980-Baz), which maintains apico-basal polarity and adherens junction assembly and positioning (Tepass, 2012; Zou et al., 2012; Roper, 2012; Letizia et al., 2013; Pichaud, 2018). In flies, the Crb transmembrane domain retains Cdc42-Par6-aPKC at the apical part of the cell, enabling PS980-Baz to accumulate at the lateral part of the cell and recruit adherens junction material, including E-cadherin, a key intercellular adhesion factor (Tepass, 2012; Pichaud, 2018). Our experiments suggest that the association of CRB2S with CRB2 disrupts this pathway. Exposure of cells to CRB2S results in the loss of aPKC, the mediator of PAR complex signalling (Goldstein and Macara, 2007; St Johnston and Ahringer, 2010; Pichaud, 2018), and disrupts E-cadherin localisation. Together our findings suggest a model (Fig.6H and Fig 7O) in which all VL cells, including dmNes^+^RG cells express CRB2, which supports apico-basal polarity; however, dmNes^+^RG also secrete CRB2S, which acts non cell-autonomously and locally to compete away CRB2 in neighboring cells, leading to loss of apico-basal polarity and delamination. Together these findings add to the evidence that the fly regulatory network is conserved and suggest a mechanism through which CRB2S can disrupt neuroepithelial VL integrity.

Static images suggest that dorsal collapse proceeds in an ordered fashion: dorsal-most progenitor cells delaminate first, then more ventral cells. Live tissue imaging confirmed this, showing that the progenitor cell that is adjacent to the dmNes^+^RG abscises and delaminates. This process continues in a progressive manner, and as each progenitor cell delaminates, the dmNes^+^RG end-foot ratchets down. What orchestrates progressive delamination and ratcheting? Previous in vivo studies in zebrafish have suggested that dmNes^+^RG cells are tethered to VL cells via a filamentous (F)-actin cytoskeleton belt, that tethering is required for dorsal collapse and depends on the activity of Rho-associated protein kinase (Rok) (Kondrychyn et al., 2013). In Drosophila, Crumbs acts via aPKC to negatively regulate Rok, which in turn prevents formation of a supracellular actomyosin cable: thus in tubulogenesis, low levels of Crumbs/aPKC correlate with formation of a cable that is under increased tension (Roper, 2012). We likewise detect a continuous F-actin belt that runs through dmNes^+^RG to VL cells, throughout collapse, and observe that the loss of CRB2 and aPKC is limited to VL progenitor cells that lie immediately adjacent to dmNes^+^RG. Therefore, one prosaic possibility is that local loss of CRB2 and aPKC promotes formation of a supracellular actin cable which ratchets dmNes^+^RG end-feet to the next VL cell and tensions the diminishing VL. Although this idea remains to be tested, our data adds to previous studies that show that dorsal collapse is mediated through intimate interactions between dmNes^+^RG and VL cells, and we propose that the local loss of CRB2/aPKC is instrumental to formation of a continuous cable, which limits the action of CRB2S to ensure integrity of the VL epithelium while individual cells delaminate. Potentially, then, a homophilic binding of CRB2S-CRB2 causes downregulation of aPKC to orchestrate both progenitor cell delamination and formation of a supracellular actin cable.

Our findings do not address how VL progenitor cell delamination initiates or terminates, but one possibility is that this is effected by the action of CRB2S over a narrow time window. Little CRB2 is detected in dmNes^+^RG at E13.5 (Fig.S3B), prior to dorsal collapse (and relatively little Crb2s-encoding mRNA is detected at E11.5). Further, the punctate, non-apical CRB2 that we detect at E15.5 (Fig. 5B,B’), potentially CRB2S, is not obviously detected at the end of dorsal collapse (Fig.S3E,F). Another possibility, not exclusive, is that ventral VL cells are, or become, refractory to the action of CRB2S. Future studies are needed to understand this question, and that of why dmNes^+^RG are themselves refractory to the action of CRB2S.

The mechanism that we uncover in these studies is likely to be one of multiple steps that contribute to spinal cord ventricular layer attrition and eventual formation of the adult central canal. The looser arrangement of dorsal VL cells and their lack of expression of Zo-1 indicates that the attrition process we describe here is only one of multiple integrated mechanisms. Previous studies, for instance, indicate a role for declining proliferation (Cañizares et al, 2019) and Wnt signalling (Xing et al., 2018). Moreover, a parallel study in one of our labs reveals that central canal formation proceeds through a combination of cell rearrangements at each pole: dorsally, dorsal collapse and ventrally, a dissociation of a subset of floor plate cells, accompanied by changes in the activity of critical ventral and dorsal patterning signals: a gradual decline in ventral sonic hedgehog activity and an expansion of dorsal bone morphogenetic protein signalling (Cañizares et al, 2019). Our data here provides evidence that CRB2 is involved in these additional events: in mice that lack CRB2, cell fate specification mediated by patterning signals is dysregulated: no expression of the homeobox transcription factor, Nkx6.1, is detected on aberrantly-retained VL progenitors. Further studies are needed to determine whether CRB2 governs Nkx6.1 through its ability to interact with other homeobox genes (Lee et al 2011), or via an effect on Shh-BMP signalling.

In conclusion, our studies reveal a novel mechanism of action of CRB2, in which the action of a secreted variant, CRB2S, deriving from dmNes^+^RG, mediates local delamination, as part of a mechanism that establishes the central canal. The finding that CRB2S is expressed elsewhere in the CNS suggests it may operate more widely to promote local delamination: future studies are needed to establish whether its expression in SVL cells of the lateral ventricle, a recognized stem cell niche in the brain, promotes delamination associated with neuronal differentiation, and whether its expression in the eye is involved in dynamism of the retinal pigment epithelium (RPE), where Crb2 mutations trigger EMT and promote RPE degeneration (Alves et al., 2013; Alves et al., 2014; Chen et al., 2019). More generally, our findings demonstrate how early patterning centres (dmNes+RG derive from roof plate cells) are maintained through life to support remodelling and maintenance, add to the evidence that, similar to *Drosophila* epithelial sheets, the vertebrate neuroepithelium is modelled by dynamic local cell-cell interactions, and reveals a cell non-autonomous action for CRB2S in neuroepithelial remodelling.

## Acknowledgements

We thank Natalia Bulgakova for help with interpretation of EM images and Andrew Furley for helpful comments

## Method

### Mice

C57black/6J or CD1 mice were used to obtain wild type mouse embryos. Timed mating was used to obtain embryos at the appropriate stages. Pregnant mice were anaesthetised to unconsciousness through isoflurane inhalation (Isoflo, Abbot) before cervical dislocation. Embryos were transferred into Leibovitz’s 15 (L-15, Gibco) before decapitation, then further dissection. All procedures concerning transgenic animals were performed with the permission of the animal experimentation committee (DEC) of the Royal Netherlands Academy of Arts and Sciences (KNAW) (permit number NIN06–46) or (for wild types) the ethical committee of The University of Sheffield.

### Generation of transgenic mice

To generate CRB2 Nestin-Cre cKO (Nestin-Cre^+/-^/CRB2^F/F^) mouse embryos, homozygote CRB2^F/F^ mice (Alves et al., 2013) were crossed at the Netherlands Institute for Neurosciences with double heterozygote Nestin-Cre^+/-^ (Dubois et al., 2006)/CRB2^F/+^ mice. Nestin-Cre expressing cells specifically delete CRB2 encoding exons 10-13 (Alves et al., 2014). Note CRB2 Nestin-Cre cKO mice die shortly after birth. For the analysis of the mutant mouse models, embryos were genotyped and sent to the UK in 30% sucrose solution. Three control and three conditional knockout embryos were used for marker analysis.

### Immunohistochemistry

Tissues were fixed in 4% paraformaldehyde for 2 hrs, transferred to 30% sucrose overnight, then cryosectioned at 15μm and incubated with primary antibodies: anti-Nestin (1:300, AbCAM), anti-Nkx6.1 (1:50, DSHB), anti-Sox1 (1:300, Cell Signalling Technologies), anti-Sox2 (1:1000, Millipore), anti-Sox3 (1:1000 Gift from T.Edlund), anti-Zo-1 (1:300, Zymed) anti-Phalloidin (1:500, Thermofisher), anti-aPKC (1:300, Santa-Cruz), anti-PAR3 (1:200, Millipore), anti-CRB2 (1:300, Boroviak and Rashbass, 2011), anti-Dystroglycan (1:30, Gift from S.Winder), anti-M2 (1:50, DSHB), anti-Transitin (1:50, DSHB), anti-Pax6 (1:50, DSHB) and anti-pH3 (1:1000, Millipore). Alexa 488 and 594 conjugated secondary antibodies were used (1:500; ThermoFisher Scientific/Molecular Probes, cat. nos. A11001, A11034 and A11005). Slides were mounted in Vectashield (Vector Laboratories) and analysed. For each antibody, three sections were analysed from 3 embryos at each stage. For analysis of laminin/dystroglycan, breaks in basement membrane were scored as regions >3 nuclei in length, to avoid counting small tears.

### RT-PCR

The Crbs2S RT-PCR reaction was performed on cDNA synthesized from tissues using SuperScript III First-Strand Synthesis System (Invitrogen). *Crb2* mRNA was amplified. Primers were designed to amplify full length mature *Crb2* and *Crbs2*. A second round of PCR was then performed, designed to amplify *Crb2S* specifically. A GAPDH loading control was run. The reactions were run on an agarose gel with the addition of ethidium bromide (Bio-Rad) and bands of the appropriate size were excised using QiAquick Gel Extraction Kit (Qiagen) and sequenced in-house.

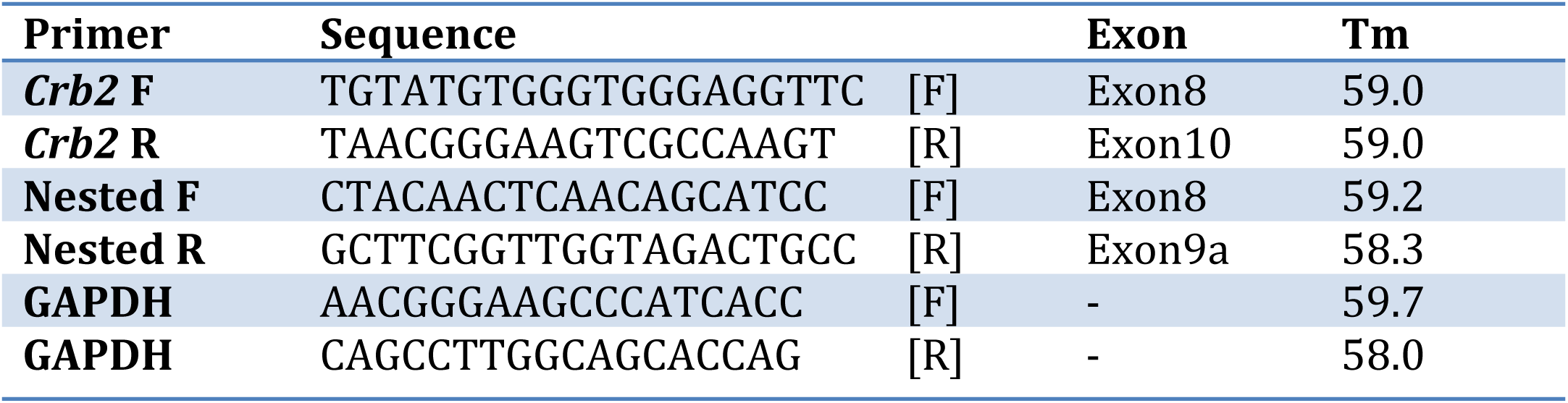

### Slice culture and live imaging

Freshly dissected E13 mouse thoracic/lumbar spinal cord was electroporated with GFP-GPI and RFP-H2B plasmids at a concentration of 0.3ug/ul. The spinal cord was mounted in low melting point agarose (Sigma) then sectioned at 300μm on a vibrating blade microtome (Leica VT1200 S). Slices were then embedded in collagen in imaging dishes (World Precision Instruments) and incubated at 37oC 50% CO2 in Neurobasal media (Gibco) supplanted with Glutamax (Gibco), B27 (Gibco), foetal calf serum (Sigma) and Gentamycin (Gibco) for 24 hours before imaging. Live imaging was performed as described in Das and Storey, 2014.

### Cell culture

Confluent MDCK cells were trypsinised and re-suspended. Cells were re-plated at low density. The cells were incubated for 6 hours and allowed to reattach. CRB2S or PBS was added to the media (see note). Cells were fixed and analysed after a 15 hour incubation.

### Light microscopy and image analysis

Fluorescent images were taken on a Zeiss Apotome 2 microscope with Axiovision software (Zeiss) or for high magnification images, on a Nikon Ti system running Nikon Elements AR software or a Deltavision RT system running SoftWorx. Timelapse images were taken on a Deltavision RT system. Images were processed using Image-J (FIJI) and made into composites using Adobe photoshop.

### Transmission electron microscopy

Specimens were fixed in 3% Glutaldehyde/0.1M Sodium Cacodylate buffer overnight, washed in buffer and dehydrated in ethanol, cleared in epoxypropane (EPP) and infiltrated in 50/50 araldite resin:EPP mixture on a rotor. This mixture was replaced twice over 8 hours with fresh araldite resin mixture before embedding and curing at 60 degrees C for 48-72 hours. Ultrathin sections, approximately 85nm thick, were cut on a Leica UC 6 ultramicrotome onto 200 mesh copper grids, stained for 30 mins with saturated aqueous Uranyl Acetate followed by Reynold’s Lead Citrate for 10 mins. Sections were examined using a FEI Tecnai Transmission Electron Microscope at an accelerating voltage of 80Kv. Electron micrographs were recorded using a Gatan Orius 1000 digital camera and Gatan Digital Micrograph software.

### *In vivo* Manipulations

Chick embryos were staged using the Hamburger-Hamilton embryo staging chart and the clear vitelline membrane was removed over the caudal neural tube into which the tissue/bead was to be transplanted. Freshly dissected embryonic mouse spinal cord was sliced into 400μm sections on a tissue chopper (McIlwain) and the slices placed into ice-cold L-15. Tissue to be transplanted was punched out with a pulled glass needle (1mm x 0.78, Harvard Apparatus) and mouth pipette (Sigma), before being carefully placed into the most rostral part of the open neural tube. Affi-gel beads (Bio-Rad) were soaked in protein or PBS control for 24 hours before transplantation into RFP electroporated/non-electroporated HH10 chick embryos.. Beads were carefully placed into the most rostral part of the open neural tube. The egg was sealed before incubation at 37°C for 24 hours. Embryos were then dissected out and fixed in ice-cold 4% PFA in for 2 hours prior to sectioning and immunohistochemistry.

### Generation of stable cell lines and CRB2S protein purification and sequencing

HEK293 cells were transfected with either CRB2S cDNA or CRB2 signal peptide cDNA expression vectors. The cells were grown and expanded in selective conditions - medium+ G418 (800 μg/ml-Sigma Aldrich). The expression level of protein of interest in the clones was determined using V5 tagged protein in the processed cell culture supernatant by Western blotting. Three positive clones were expanded.

A HEK 293 stable cell line overexpressing *Crb2S* was used for obtaining purified protein. The transgenic *Crb2S* cell line was passed onto BioServ UK for scale-up of cells and immobilized metal ion affinity chromatography (IMAC). The cells were maintained in G418 selection antibiotic (800 μg/ml) throughout the culture period. The purified CRB2S protein (100 μg/ml) was sequenced as described below, aliquoted and stored at −80°C. For protein sequencing, SDS gel electrophoresis was carried out as described below; care was taken to minimize external keratin contamination from the environment. All processing was carried out in a clean biosafety cabinet. The gel was fixed and stained with Coomassie Brilliant Blue (Sigma-Aldrich) as per manufacturer’s instructions and the bands of interest were excised using a clean blade and stored at 4°C in a sterile tube. LC-ESI-Mass spectrometry was carried out by a commercial company (Eurogentec) using an LC (nano-Ultimate 3000- Dionex)-ESIion trap (AMAZONE-Bruker) in positive mode.

### Western Blotting

Western blotting was carried out using the NuPAGE gel system (Invitrogen), according to the manufacturer’s instructions. Cells were lysed in RIPA buffer (Upstate) supplemented with protease inhibitor tablets (complete Mini, EDTA free, Roche) on ice. 20μg total protein lysate, as determined by Bradford assay, per sample were loaded on a 4-12% gradient gel (Invitrogen) under denaturing conditions. Membranes (Hybond-C Extra, Amersham Biosciences) were blocked in 5% semi skimmed milk powder, Tween (0.1%) PBS and primary antibodies (V5 Tag Abcam, chicken polyclonal 1:2000, His Tag Cell signalling, rabbit polyclonal 1:1000) were applied in blocking solution overnight at 4°C. Secondary antibodies linked to horse radish peroxidase (all from Stratech) were applied for 1h at a concentration of 1:1000.

## References

Alaynick WA, Jessell TM and Pfaff SL. (2011). SnapShot: spinal cord development. Cell. 2011 Jul 8;146(1):178–178.e1. doi: 10.1016/j.cell.2011.06.038.

Altman J and Bayer S.A. (1984) The development of the rat spinal cord. Springer-Verlag, Berlin.

Alves CH, Sanz AS, Park B, Pellissier LP, Tanimoto N, Beck SC, Huber G, Murtaza M, Richard F, Sridevi Gurubaran I, Garcia Garrido M, Levelt CN, Rashbass P, Le Bivic A, Seeliger MW, Wijnholds J. (2013) Loss of CRB2 in the mouse retina mimics human retinitis pigmentosa due to mutations in the CRB1 gene. Hum Mol 22(1): 35–50. doi: 10.1093/hmg/dds398. PubMed PMID: 23001562.

Alves CH, Pellissier LP, Vos RM, Garcia Garrido M, Sothilingam V, Seide C, Beck SC, Klooster J, Furukawa T, Flannery JG, Verhaagen J, Seeliger MW, Wijnholds J. (2014) Targeted ablation of CRB2 in photoreceptor cells induces retinitis pigmentosa. Hum Mol Genet. 23(13):3384–401. doi: 10.1093/hmg/ddu048.

Assemat, E., Bazellieres, E., E. Pallesi-Pocachard, E., A. Le Bivic, A. and Massey-Harroche, D. (2008) Polarity complex proteins Biochem. Biophys. Acta, 1778 pp. 614–630

Bazellières E, Aksenova V, Barthélémy-Requin M, Massey-Harroche D, Le Bivic A. (2018) Role of the Crumbs proteins in ciliogenesis, cell migration and actin organization.SeminCellDevBiol.2018 Sep;81:13–20.doi: 10.1016/j.semcdb.2017.10.018.

Boroviak T, Rashbass P. (2011) The apical polarity determinant Crumbs 2 is a novel regulator of ESC-derived neural progenitors. Stem Cells. 29(2):193–205. doi: 10.1002/stem.567.

Bohme, G. (1988). Formation of the central canal and dorsal glial septum in the spinal cord of the domestic cat. Journal of Anatomy 159, 37–47.

Bruni, J.E. (1998). Ependymal development, proliferation, and functions: a review. Microsc Res Tech 41, 2–13.

Bulgakova, N.A. and Knust, E. (2009) The Crumbs complex: from epithelial-cell polarity to retinal degeneration. J. Cell Sci., 122, pp. 2587–2596

Cañizares, M., Rodrigo Albors, A., Singer, G., Suttie, N., Gorkic, M., Felts, P. and Storey, K.G. (2019). Multiple steps mediate ventricular layer attrition to form the adult mouse spinal cord central canal. Biorixv.

Cearns MD, Escuin S, Alexandre P, Greene ND and Copp AJ(2016). Microtubules, polarity and vertebrate neural tube morphogenesis. J Anat. 229(1):63–74.

Chen X, Jiang C, Yang D, Sun R, Wang M, Sun H, Xu M, Zhou L, Chen M, Xie P, Yan B, Liu Q, Zhao C. (2019). CRB2 mutation causes autosomal recessive retinitis pigmentosa. Exp Eye Res. 180:164–173. doi: 10.1016/j.exer.2018.12.018.

Cohen D, Brennwald PJ, Rodriguez-Boulan E, Müsch A. (2004). Mammalian PAR-1 determines epithelial lumen polarity by organizing the microtubule cytoskeleton. J Cell Biol. 164(5):717–27.

Das R. and Storey K.G. (2014) Apical Abscission Alters Cell Polarity and Dismantles the Primary Cilium During Neurogenesis, Science 10:6167 200–204.

Del Bigio, M.R. (1995). The ependyma: a protective barrier between brain and cerebrospinal fluid. Glia 14, 1–13.

Deneen, B., Ho, R., Lukaszewicz, A., Hochstim, C.J., Gronostajski, R.M., Anderson, D.J., 2006. The transcription factor NFIA controls the onset of gliogenesis in the developing spinal cord. Neuron 52, 953–680 968.

Dubois NC, Hofmann D, Kaloulis K, Bishop JM, Trumpp A (2006) Nestin-Cre transgenic mouse line Nes-Cre1 mediates highly efficient Cre/IoxP mediated recombination in the nervous system, kidney, and somite-derived tissues. Genesis 44: 355–360. DOI 10.1002/dvg.20226

Dudok JJ, Murtaza M, Henrique Alves C, Rashbass P, Wijnholds J. (2016) Crumbs 2 prevents cortical abnormalities in mouse dorsal telencephalon. Neurosci Res. 2016 Jul;108:12–23. doi: 10.1016/j.neures.2016.01.001.

Fu, H., Qi, Y., Tan, M., Cai, J., Hu, X., Liu, Z., Jensen, J., Qiu, M. (2003). Molecular mapping of the origin of postnatal spinal cord ependymal cells: evidence that adult ependymal cells are derived from Nkx6.1+ ventral neural progenitor cells. The Journal of Comparative Neurology 456, 237–244.

Goldstein, B. and Macara, IG. (2007) The PAR proteins: fundamental players in animal cell polarization. Developmental Cell 13(5):609–22.

Gouti, M., Metzis, V and Briscoe, J. (2015) The route to spinal cord cell types: a tale of signals and switches. Trend Genet. 31(6), 282–289.

Goldstein B, Macara IG. (2007) The PAR proteins: fundamental players in animal cell polarization. Dev Cell. 13(5):609–22.

Ghazale, H., Ripoll, C., Leventoux, N., Jacob, L., Azar, S., Mamaeva, D., Glasson, Y., Calvo, C.F., Thomas, J.L., Meneceur, S., Lallemand, Y., Rigau, V., Perrin, F.E., Noristani, H.N., Rocamonde, B., Huillard, E., Bauchet, L., Hugnot, J.P. (2019). RNA Profiling of the Human and Mouse Spinal Cord Stem Cell Niches Reveals an Embryonic-like Regionalization with MSX1(+) Roof-Plate-Derived Cells. Stem Cell Reports 12, 1159–1177.

Hildebrand JD, Soriano P (1999) Shroom, a PDZ domain-containing actin binding protein, is required for neural tube morphogenesis in mice. Cell 99: 485–497. doi: 10.1016/s0092-8674(00)81537-8

Hochstim C, Deneen B, Lukaszewicz A, Zhou Q, Anderson DJ. (2008) Identification of positionally distinct astrocyte subtypes whose identities are specified by a homeodomain code. Cell 133, 510–22. doi: 10.1016/j.cell.2008.02.046.

Hsu YC, Jensen AM. (2010) Multiple domains in the Crumbs Homolog 2a (CRB2a) protein are required for regulating rod photoreceptor size. BMC Cell Biol. 11:60. doi: 10.1186/1471-2121-11-60.

Jessell, TM (2000). Neuronal specification in the spinal cord: inductive signals and transcriptional cords. Nat Rev Genet 1, 20–29.

Jiménez-Amilburu, V. and Stainier D.Y.R. (2019) The transmembrane protein Crb2a regulates cardiomyocyte apicobasal polarity and adhesion in zebrafish. Development. 2019 May 7;146(9). pii: dev171207. doi: 10.1242/dev.171207.

Johansson, C.B., Momma, S., Clarke, D.L., Risling, M., Lendahl, U., Frisen, J. (1999). Identification of a neural stem cell in the adult mammalian central nervous system. Cell 96, 25–34.

Kasioulis I, Das R.M. and Storey K.G (2017). Inter-dependent apical microtubule and actin dynamics orchestrate centrosome retention and neuronal delamination. Elife. 6. pii: e26215. doi: 10.7554/eLife.26215.

Katoh, M., and Katoh, M. (2004). Identification and characterization of Crumbs homolog 2 gene at human chromosome 9q33.3. Int J Oncol, 24(3), 743–749.

Klebes, A., and Knust, E. (2000). A conserved motif in Crumbs is required for E-cadherin localisation and *zonula adherens* formation in *Drosophila*. Curr. Biol. 10, 76–85.

Kondrychyn, I., Teh, C., Sin, M., Korzh, V. (2013). Stretching morphogenesis of the roof plate and formation of the central canal. PloS One 8, e56219.

Korzh V. (2014). Stretching cell morphogenesis during late neurulation and mild neural tube defects. Dev Growth Differ. 56(6):425–33.

Laug, D., Glasgow, S.M., Deneen, B., 2018. A glial blueprint for gliomagenesis. Nature Reviews Neuroscience 9, 393–403.

Le Dreau, G and Marti, E. (2012) Dorso-ventral patterning of the neural tube: a tale of three signals. Dev Neurobiol 72, 1471–1481.

Lee K.J. and Jessell TM (1999) The specification of dorsal cell fates in the vertebrate central nervous system.Annu Rev Neurosci 22: 261–294

Lee AP, Brenner S, Venkatesh B. (2011) Mouse transgenesis identifies conserved functional enhancers and cis-regulatory motif in the vertebrate LIM homeobox gene Lhx2 locus. PLoS One. 6(5):e20088. doi: 10.1371/journal.pone.0020088.

Letizia A, Ricardo S, Moussian B, Martín N, Llimargas M. (2013) A functional role of the extracellular domain of Crumbs in cell architecture and apicobasal polarity. J Cell Sci. 126: 2157–63.

Li, X., Floriddia, E.M., Toskas, K., Fernandes, K.J., Guerout, N., Barnabe-Heider, F., (2016). Regenerative Potential of Ependymal Cells for Spinal Cord Injuries Over Time. EBioMedicine 13, 55–65.

McCaffrey LM, Macara IG. (2011) Epithelial organization, cell polarity and tumorigenesis. Trends Cell Biol. 21(12):727–35.

Marichal N, Reali C, Trujillo-Cenóz O, Russo RE. (2017). Spinal Cord Stem Cells In Their Microenvironment: The Ependyma as a Stem Cell Niche. Adv Exp Med Biol. 1041, 55–79.

Meletis, K., Barnabe-Heider, F., Carlen, M., Evergren, E., Tomilin, N., Shupliakov, O., Frisen, J. (2008). Spinal cord injury reveals multilineage differentiation of ependymal cells. PLoS Biology 6, e182.

Munson C, Huisken J., Bit-Avragim N., Kuo T, Dong PD, Ober EA, Verkade H, Abdelilah-Seyfried S.and Stainier DY.(2008).Regulation of neurocoel morphogenesis by Pard6 gamma b. Dev Biol. 324(1):41–54.

Nagele, R.G., Hunter, E., Bush, K. and Lee, H.Y. (1987) Studies on the mechanisms of neurulation in the chick: morphometric analysis of force distribution within the neuroepithelium during neural tube formation. J. Exp. Zool. 1987 Dec;244(3):425–36.

Nikolopoulou E, Galea GL, Rolo A, Greene ND and Copp AJ (2017). Neural tube closure: cellular, molecular and biomechanical mechanisms. Development. 2017 Feb 15;144(4):552–566. doi: 10.1242/dev.145904.

Omori Y, Malicki J. (2006) oko meduzy and related crumbs genes are determinants of apical cell features in the vertebrate embryo. Curr Biol. 16:945–957.

Paniagu, A.E., Herranz-Martin, S., Jimeno, D, Jimeno, A.M., Lopez-Benitro, S., Arevalo, J.C., Velasco, A., Aijon, J and Lillo, C. (2015) CRB2 completes a fully expressed Crumbs complex in the Retinal Pigment Epithelium. Sci Rep.5: 14504.

Pevny L. and Placzek M. (2005) SOX genes and neural progenitor identity. Curr Opin Neurobiol. 2005 Feb;15(1):7–13.

Pichaud F. (2018) PAR-Complex and Crumbs Function During Photoreceptor Morphogenesis and Retinal Degeneration. Front Cell Neurosci. 12:90. doi: 10.3389/fncel.2018.00090.

Ramkumar N, Harvey BM, Lee JD, Alcorn HL, Silva-Gagliardi NF, McGlade CJ, Bestor TH,Wijnholds J, Haltiwanger RS, Anderson KV.(2015)ProteinO-Glucosyltransferase 1 (POGLUT1) Promotes Mouse Gastrulation through Modification of the Apical Polarity Protein CRUMBS2. PLoS Genet. 11:e1005551.

Ramkumar N, Omelchenko T, Silva-Gagliardi NF, McGlade CJ, Wijnholds J and Anderson KV. (2016) Crumbs2 promotes cell ingression during the epithelial-to-mesenchymal transition at gastrulation. Nat Cell Biol. 18(12):1281–1291. doi: 10.1038/ncb3442. Epub 2016 Nov 21.

Richard M, Roepman R, Aartsen WM, van Rossum AG, den Hollander AI, Knust E, Wijnholds J, Cremers FP. (2006) Towards understanding CRUMBS function in retinal dystrophies. Hum Mol Genet. 15 Spec No 2:R235–43.

Roper, K. (2012). Anisotropy of Crumbs and aPKC drives myosin cable assembly during tube formation. Dev. Cell 23, 939–953. doi: 10.1016/j.devcel.2012.09.013

Sabourin, J.C., Ackema, K.B., Ohayon, D., Guichet, P.O., Perrin, F.E., Garces, A., Ripoll, C., Charite, J., Simonneau, L., Kettenmann, H., Zine, A., Privat, A., Valmier, J., Pattyn, A., Hugnot, J.P. (2009). A mesenchymal- 780 like ZEB1(+) niche harbors dorsal radial glial fibrillary acidic protein-positive stem cells in the spinal cord. Stem Cells 27, 2722–2733.

Sadler TW, Greenberg D, Coughlin P, Lessard JL (1982) Actin distribution patterns in the mouse neural tube during neurulation. Science 215: 172–174. doi: 10.1126/science.7031898

Sawyer JM, Harrell JR, Shemer G, Sullivan-Brown J, Roh-Johnson M, Goldstein B. (2010) Apical Constriction: A Cell Shape Change that Can Drive Morphogenesis. Dev Biology. 341:5–19. doi:10.1016/j.ydbio.2009.09.009.

Sevc, J., Daxnerova, Z., Miklosova, M. (2009). Role of radial glia in transformation of the primitive lumen to 788 the central canal in the developing rat spinal cord. Cell Mol Neurobiol 29, 927–936.

Shinozuka T, Takada R, Yoshida S, Yonemura S and Takada S (2019) Wnt produced by stretched roof-plate cells is required for the promotion of cell proliferation around the central canal of the spinal cord Development 2019 146: dev159343 doi: 10.1242/dev.159343

Snow. D, M., Steindler, D, A., Silver, J (1990) Molecular and cellular characterization of the glial roof plate of the spinal cord and optic tectum: A possible role for a proteoglycan in the development of an axon barrier. Developmental Biology, Volume 138, issue 2, Pages 359–376

Spassky N, et al. (2005) Adult ependymal cells are postmitotic and are derived from radial glial cells during embryogenesis. J Neurosci. 2005;25:10–18.

St Johnston D. and Ahringer, J. (2010) Cell polarity in eggs and epithelia: parallels and diversity. Cell 141(5):757–74. doi: 10.1016/j.cell.2010.05.011.

Stolt, C.C., Lommes, P., Sock, E., Chaboissier, M.C., Schedl, A., Wegner, M., 2003. The Sox9 transcription factor determines glial fate choice in the developing spinal cord. Genes & Development 17, 1677–1689.

Sturrock, R.R. (1981). An electron microscopic study of the development of the ependyma of the central canal of the mouse spinal cord. Journal of Anatomy 132, 119–136.

Tepass, U. (2012) The Apical Polarity Protein Network in *Drosophila*Epithelial Cells: Regulation of Polarity, Junctions, Morphogenesis, Cell Growth, and Survival. Annu Rev Cell Dev Biol. 2012;28:655–85.

Ulloa, F. and Briscoe, J. (2007) Morphogens and the control of cell proliferation and patterning in the spinal cord. Cell Cycle 6, 2640–2649.

van de Pavert SA, Kantardzhieva A, Malysheva A, Meuleman J, Versteeg I, Levelt C, Klooster J, Geiger S, Seeliger MW, Rashbass P, Le Bivic A, Wijnholds J (2004) Crumbs homologue 1 is required for maintenance of photoreceptor cell polarization and adhesion during light exposure. J Cell Sci. 117(Pt 18):4169–77.

Van den Hurk J.A., Rashbass P., Roepman R., Davis J., Voesenek K.E., Arends M.L., Zonneveld M.N., van Roekel M.H., Cameron K., Rohrschneider K, et al. (2005) Characterization of the Crumbs homolog 2 (*CRB2*) gene and analysis of its role in retinitis pigmentosa and Leber congenital amaurosis, Mol. Vision 11 263–273

Wakamatsu Y, Nakamura N, Lee I-A, Cole GJ, Osumi N (2007) Transitin, a nestin-like intermediate filament protein, mediates cortical localization and the lateral transport of Numb in mitotic avian neuroepithelial cells. Development 134: 2425–2433; doi: 10.1242/dev.02862

Watanabe, T., Miyatani, S., Katoh, I., Kobayashi, S., & Ikawa, Y. (2004). Expression of a novel secretory form (Crb1s) of mouse Crumbs homologue Crb1 in skin development. Biochemical and Biophysical Research Communications, 313(2), 263–270.

Whalley K, Gögel S, Lange S, Ferretti P. (2009). Changes in progenitor populations and ongoing neurogenesis in the regenerating chick spinal cord. Dev Biol. 2009 Aug 15;332(2):234–45. doi: 10.1016/j.ydbio.2009.05.569.

Wodarz A, Hinz U, Engelbert M, Knust E. (1995) Expression of crumbs confers apical character on plasma membrane domains of ectodermal epithelia of Drosophila. Cell 82(1):67–76.

Xiao Z, Patrakka J, Nukui M, Chi L, Niu D, Betsholtz C, Pikkarainen T, Vainio S, Tryggvason K. (2011) Deficiency in Crumbs homolog 2 (CRB2) affects gastrulation and results in embryonic lethality in mice. Dev Dyn. 240(12):2646–56. doi: 10.1002/dvdy.22778.

Xing L, Anbarchian T, Tsai JM, Plant GW and Nusse R (2018). Wnt/β-catenin signaling regulates ependymal cell development and adult homeostasis. Proc Natl Acad Sci U S A. 115(26):E5954–E5962. doi: 10.1073/pnas.1803297115.

Yu K, McGlynn S, Matise MP. Floor plate-derived sonic hedgehog regulates glial and ependymal cell fates in the developing spinal cord. Development. 2013 Apr;140(7):1594–604. doi: 10.1242/dev.090845.

Zou J, Wang X, Wei X. (2012) Crb apical polarity proteins maintain zebrafish retinal cone mosaics via intercellular binding of their extracellular domains. Dev Cell. 22(6):1261–74. doi: 10.1016/j.devcel.2012.03.007.

